# TL1A/DR3 signaling regulates the generation of pathogenic Th9 cells in experimental inflammatory bowel disease

**DOI:** 10.1101/2024.02.09.579684

**Authors:** Paola Menghini, Ludovica F. Buttó, Adrian Gomez-Nguyen, Natalia Aladyshkina, Kristine-Ann Buela, Abdullah Osme, Ricky Chan, Giorgos Bamias, Theresa Pizarro, Fabio Cominelli

## Abstract

**Objective:** Death receptor 3 (DR3) and its ligand tumor necrosis factor like ligand 1A (TL1A), are involved in the regulation of the balance between effector and regulatory T cells in IBD. New evidence suggests a role of IL-9-secreting Th9 cells in the pathogenesis of ulcerative colitis (UC), although the molecular pathways through which IL-9 and Th9 cells may mediate intestinal inflammation in Crohn’s disease (CD) are still unclear.

**Design:** We investigated the role of DR3 signaling in the differentiation of Th9 cells in mouse models of CD-like ileitis and colitis, including SAMP1/YitFc (SAMP) mice.

**Results:** Polarized-Th9 cells with functional DR3 from SAMP WT (Th9_WT_) harbor a pro-inflammatory signature compared to DR3-deficient Th9 cells that were obtained from DR3^-/-^×SAMP mice (Th9_KO_). Conversely, ablation of DR3 signaling generated anti-inflammatory responses, as reflected by higher numbers of IL-10 producing cells in DR3^-/-^×SAMP mice. Additionally, RNA-seq and phosphoproteomic analyses showed that inflammatory pathways are significantly more activated in Th9_WT_ than in Th9_KO_ cells. Finally, in the T-cell adoptive transfer model, Th9_KO_ cells were less colitogenic than Th9_WT_, while IL-9 blockade diminished the severity of intestinal inflammation, indicating a crucial role of functional DR3 receptor in Th9 cells pathogenicity.

**Conclusion:** We describe herein that a functional DR3 receptor is required for the pathogenicity of Th9 cells, thus, constituting a novel mechanism by which TL1A/DR3 signaling mediates experimental CD-like ileitis. The TL1A/DR3/Th9 pro-inflammatory pathway may offer a novel therapeutic target for patients with CD.

**What is already known on this topic**

– TL1A/DR3 system plays a pivotal role in the pathogenesis of Crohn’s disease like ileitis.
– Th9 cells are a novel subset of T lymphocytes mainly producing the pro-inflammatory cytokine IL-9 which contributes to intestinal inflammation.
– Those finding provided us with a strong rationale to investigate IL-9-producing cells in our SAMP1/YitFc mouse model of CD-like ileitis.

**What this study adds**

– DR3 receptor is involved in the regulation and progression of intestinal inflammation by promoting Th9 cell differentiation and pathogenicity.
– Using RNA-seq and phosphoproteomic comparative analyses we were able to characterize Th9 cells with and without functional DR3 receptor showing that presence of DR3 confers a pro-inflammatory signature to Th9 cells.
– We describe a novel role of DR3 in Th9 cells development that appears to regulate their pro-inflammatory phenotype in models of CD-like ileitis and colitis.

**How this study might affect research, practice or policy**

– TLA1/DR3 axis and Th9 cells may be useful as therapeutic targets in CD.

## Introduction

Crohn’s disease (CD) and ulcerative colitis (UC) are the two dominant phenotypes of inflammatory bowel diseases (IBD), the hallmark of which is chronic, remitting and relapsing inflammation of the intestinal tract^1^. Despite recent advances in medical therapy for CD, a significant number of patients do not respond initially or lose response overtime and may experience adverse side effects that lead to treatment cessation. As a result, CD imposes a major health burden worldwide with a significant proportion of patients requiring surgical interventions^2, 3^. These facts point out a significant unmet need in the treatment for CD and emphasize the need for alternative therapeutic targets.

The current dogma for CD pathogenesis implicates dysregulated T helper (Th) cell responses and pro-inflammatory cytokines as critical mediators of tissue damage. Previous studies on mouse models of IBD have substantiated pivotal roles for Th1, Th2, and Th17 cell subsets in triggering and perpetuating experimental chronic intestinal inflammation^4^. Recently, the repertoire of pathogenic T-helper lymphocytic subsets has been expanded to include Th9 cells, which predominantly secrete proinflammatory interleukin-9 (IL-9) and have been attributed a significant role in immune-mediated inflammatory diseases, including IBD^5, 6^. Patients with UC have high colonic and systemic amounts of IL-9 as compared to healthy controls^7^. Additionally, IL-9 blockade ameliorated disease severity in experimental models of intestinal inflammation, thus, suggesting that targeting IL-9 may represent a novel therapeutic approach for UC^8, 9^. Nevertheless, both pro- and anti-inflammatory properties have been demonstrated for Th9 cells, thus, suggesting potentially dichotomous roles depending on the specific scenario (homeostasis vs. inflammation)^10^. Moreover, the regulation and downstream effectors of the TH9/IL-9 pathway during intestinal inflammation are unclear at present, as is also its potential involvement in the pathogenesis of CD.

Death receptor 3 (DR3, TNFRSF25), a member of the tumor necrosis factor receptor (TNFR) superfamily, is preferentially expressed on activated T cells and is the functional receptor for tumor necrosis factor-like cytokine 1A (TL1A, TNFSF15)^11^. The latter is expressed by dendritic cells, monocytes, macrophages, plasma cells, synovial fibroblasts, and endothelial cells. TL1A/DR3 signaling has been implicated in regulating Th1, Th2 and Th17 effector, as well as regulatory T-cell responses and innate lymphoid cell functions during immune-mediated diseases^12^. In recent years, several lines of clinical and experimental evidence have implicated the TL1A/DR3 system in the pathogenesis of IBD. Specifically, TL1A and DR3 expression is significantly increased in both serum and inflamed tissues in IBD patients and in murine experimental ileitis^13^. Sustained expression of TL1A leads to chronic intestinal inflammation in mice, whereas blockade of the TL1A/DR3 axis suppresses murine colitis^14^. Our laboratory has recently shown that TL1A/DR3 is a key regulator of experimental ileitis and that deletion of DR3 in ileitis-prone SAMP1/YitFc (SAMP) mice, leads to downregulation of pro-inflammatory genes and imbalance between Teff and Treg in the small intestine^15^. These experimental data set the background for the development of pharmacological agents that target the TL1A/DR3 axis, and which are currently tested in clinical trials in patients with IBD^16^.

In the present study, we hypothesized that DR3 is involved in the development of CD-like intestinal inflammation by promoting the differentiation of Th9 cells. To test our hypothesis, we investigated the differences between Th9 cell subsets obtained from ileitis-prone SAMP mice that were either wild-type (Th9_WT_) or had a deletion for *Dr3* [DR3^-/-^×SAMP (Th9_KO_)]. Our results show that a functional DR3 receptor significantly contributes to the activation of downstream signaling pathways relevant to Th9 cell differentiation and pathogenicity in SAMP mice, including proinflammatory cytokine and chemokine genes, which are critically involved in signaling pathways of IBD, Th1, and Th2 cell differentiation. Phosphoproteomic comparative analysis further confirmed that inflammation-associated pathways are significantly upregulated in Th9_WT_ than in Th9_KO_ cells and provided novel phosphoproteomic markers that can be further investigated to validate their potential use as novel molecular targets in the treatment of CD. Finally, we provide mechanistic evidence by showing that Th9_KO_ cells are less colitogenic than Th9_WT_, while IL-9 blockade diminishes the severity of intestinal inflammation in the T cell adoptive transfer model, indicating a crucial role of functional DR3 receptor in Th9 cells pathogenic potential. Collectively, our results describe a novel mechanism by which TL1A/DR3 signaling mediates CD-like ileitis that can be targeted for the treatment of CD.

## Results

### TL1A/DR3 signaling positively regulates the development and pro-inflammatory potential of Th9 cells

In the current study we utilized the SAMP1/YitFc mouse, which spontaneously develops terminal ileitis that is reminiscent of CD in several clinicopathological characteristics^17^. Thus, the generation of a SAMP1 strain with ablation of the *Dr3* gene (DR3^-/-^×SAMP) allowed us to examine the effect of DR3 signaling on the development of Th9 cells in relation to the pathogenesis of SAMP1 ileitis. As previously shown^15^, DR3^-/-^×SAMP (KO) mice exhibited significantly reduced intestinal inflammation compared to their wild-type (WT) counterparts, as depicted in 3D-stereomicroscopy analysis of fixed postmortem ileal specimens (Fig. S1A, 1B) and histological grading (Fig. S1C, 1D) ^11, 15^. Next, we focused on the effect of DR3 deletion on IL-9 production and the expression of genes that encode for specific proteins that affect the Th9 lineage. Our analysis showed that, as compared to WT mice, ileal samples from SAMP1 KO exhibited down-regulated expression for *il-9* and *batf3* (a positive regulator for Th9 differentiation)^18, 19^ and, inversely, elevated expression for several IL-9 inhibitory genes (*Gata3*, *Tak1*, *Id3*), (Fig. 1A). Accordingly, ileal homogenates from SAMP KO mice were significantly depleted from IL-9 protein, as compared to WT mice (Fig. 1B). Notably, Pearson correlation analysis revealed a highly positive association between IL-9 content and total inflammatory score in SAMP WT (Fig. 1C). These findings suggest that functional DR3 signaling is necessary for optimal IL-9 pathway activation.

**Figure 1:**
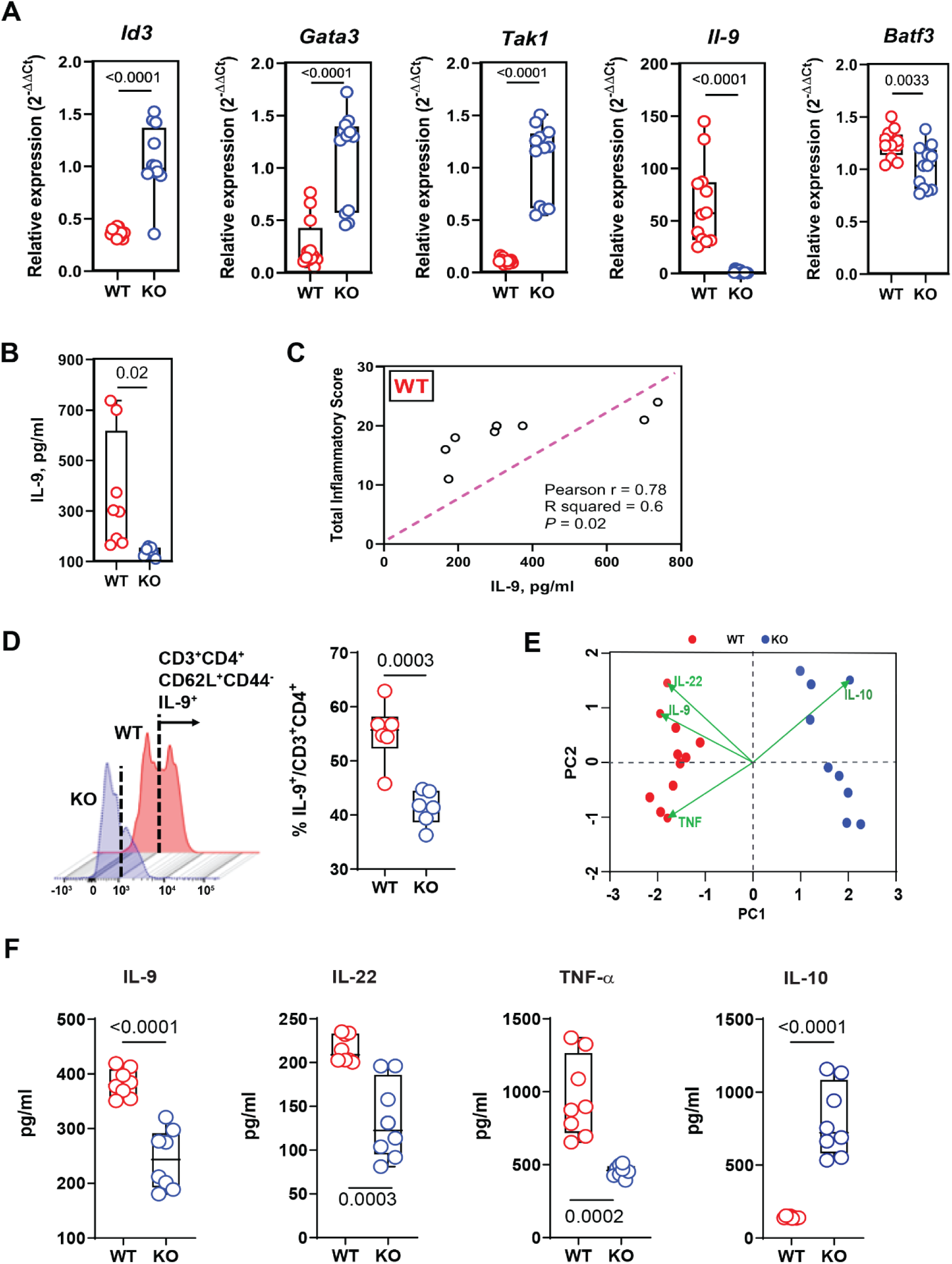
DR3 deletion reduces IL-9 levels in SAMP mice. (**A**) Relative expression of indicated gene mRNAs measured in total tissue from the ileum of WT and KO mice. The mRNA levels were determined by RT-qPCR, normalized to β-actin and expressed as fold change (2^−ΔΔCt^) (n=10). (**B**) IL-9 protein measured in tissue homogenates from the ileum of WT and KO mice. (**C**) Pearson’s correlation between IL-9 concentration and inflammatory score calculated in WT (n=8). (**D**) Intracellular IL-9 production detected by flow cytometry in MLN cells from 20-wk old SAMP and SAMPxDR3^-/-^ mice incubated for 3 days under Th9 polarizing condition supplemented with TL1A ligand, followed by stimulation with PMA, ionomycin and brefeldin A. (**F**) PCA of cytokines concentration measured in supernatants of TH9 polarized cells from WT and KO mice and (**G**) bar chart displaying their single values (n=9). Data, indicated as mean ± SD, correspond to 2 independent experiments; in (A), (B), (D), (G) statistical analysis was determined by 2-tailed Unpaired t-test.

The data above provided us with the rationale to investigate IL-9-producing cells in the SAMP1 model of CD-like ileitis. To this end, we obtained naïve CD4^+^ T cells (identified as CD4^+^, CD44^lo^, CD62L^hi^, and CD25^lo^) from the spleens of 20-wk-old AKR (uninflamed control), SAMP WT, and DR3^-/-^×SAMP mice and cultured them under Th9-polarizing conditions (i.e. TGF-β plus IL-4), with or without addition of TL1A (Fig. S2). We first observed that TL1A is essential for Th9 differentiation, as much as a functional DR3 receptor. In fact, we were able to differentiate Th9 cells from FACS sorted CD4^+^ naïve T cells only in the presence of a functional TL1A/DR3 axis (Fig. S3). We observed that the presence of TL1A in the culture medium led to significantly increased concentration of IL-9 protein in the culture supernatants and that this effect was specific for cells isolated from SAMP WT mice as compared to cells from DR3^-/-^×SAMP or AKR mice (Fig. S3). This was further confirmed by flow cytometry which showed a significantly larger percentage of IL-9 producing cells in SAMP WT than KO mice (Fig. 1D). We next compared cytokine production of polarized, TL1A-stimulatedTh9 cells obtained from WT and KO SAMP mice (indicated respectively as Th9_WT_ and Th9_KO_). Principal component analysis of selected cytokines shows that Th9_WT_ cells have a pro-inflammatory signature characterized by higher secretion of inflammatory cytokines including IL-9, IL-22, and TNF compared to Th9_KO_ cells (Figs. 1E, 1F). In contrast, Th9_KO_ cells displayed significantly higher levels of the anti-inflammatory IL-10 protein, (Figs. 1E, 1F). Taken together, these findings support a pro-inflammatory role for the functional DR3 receptor in polarized Th9 cells.

### TL1A/DR3 signaling induces differential transcriptomic profiles on Th9 cells

We next used a high-throughput RNA-sequencing approach to explore the transcriptional differences in polarized-Th9 cells from SAMP mice with and without functional DR3 receptor. Our analysis identified 741 differentially expressed genes (DEGs, upregulated: 437; downregulated: 315) in Th9_KO_ compared to Th9_WT_ out of a total of 17,633 tested genes (Fig. 2A, B). Heatmaps shown in Fig. 2C and D depict the 50 most up- and down-regulated genes. Notably, in Th9_KO_ cells we found reduced expression of *CD70* and *Tnfsf4*, which are both involved in T cell mediated inflammation^20,21^ and of chemokines *CCL24* and *CCL17* that mediate intestinal inflammation^22, 23^ (Fig 2E). Additionally, gene expression for potent pro-inflammatory cytokines (*il-1α*, *il-12a*, *il-3*, and *il-5*) was also reduced in the Th9_KO_ transcriptome (Fig. 2E). To further explore the biological functions of the identified DEGs, Pathway enrichment analysis was performed using the Kyoto Encyclopedia of genes and Genome database (KEGG). KEGG pathway analysis uncovered a total of 38 pathways enriched in Th9_KO_ cells, including IBD, JAK/STAT signaling, Chemokine signaling, Cytokine-Cytokine Receptor Interaction, PI3K-AKT signaling and T Cell Receptor signaling pathways (Table 1). Network analyses generated with central nodes of those specific pathways show differentially expressed genes with log 2FC>1 and highlight their interconnections (Fig 2G). When focusing on DEGs that are present in the IBD pathway (Fig 2F), we found that inflammation-relevant genes are downregulated in Th9_KO_ cells. For instance, Th9_KO_ cells displayed decreased expression of members of the anti-apoptotic *BCL2* family; the latter are reported to be enriched in lamina propria lymphocytes from inflamed areas of CD patients and may be responsible for resistance of T cells to apoptotic signals^24^. Additionally, *CFS2*, which has been implicated in various inflammatory processes and considered a novel therapeutic target in other inflammatory conditions such as rheumatoid arthritis^25^ are also downregulated in Th9_KO_ cells.

**Figure 2:**
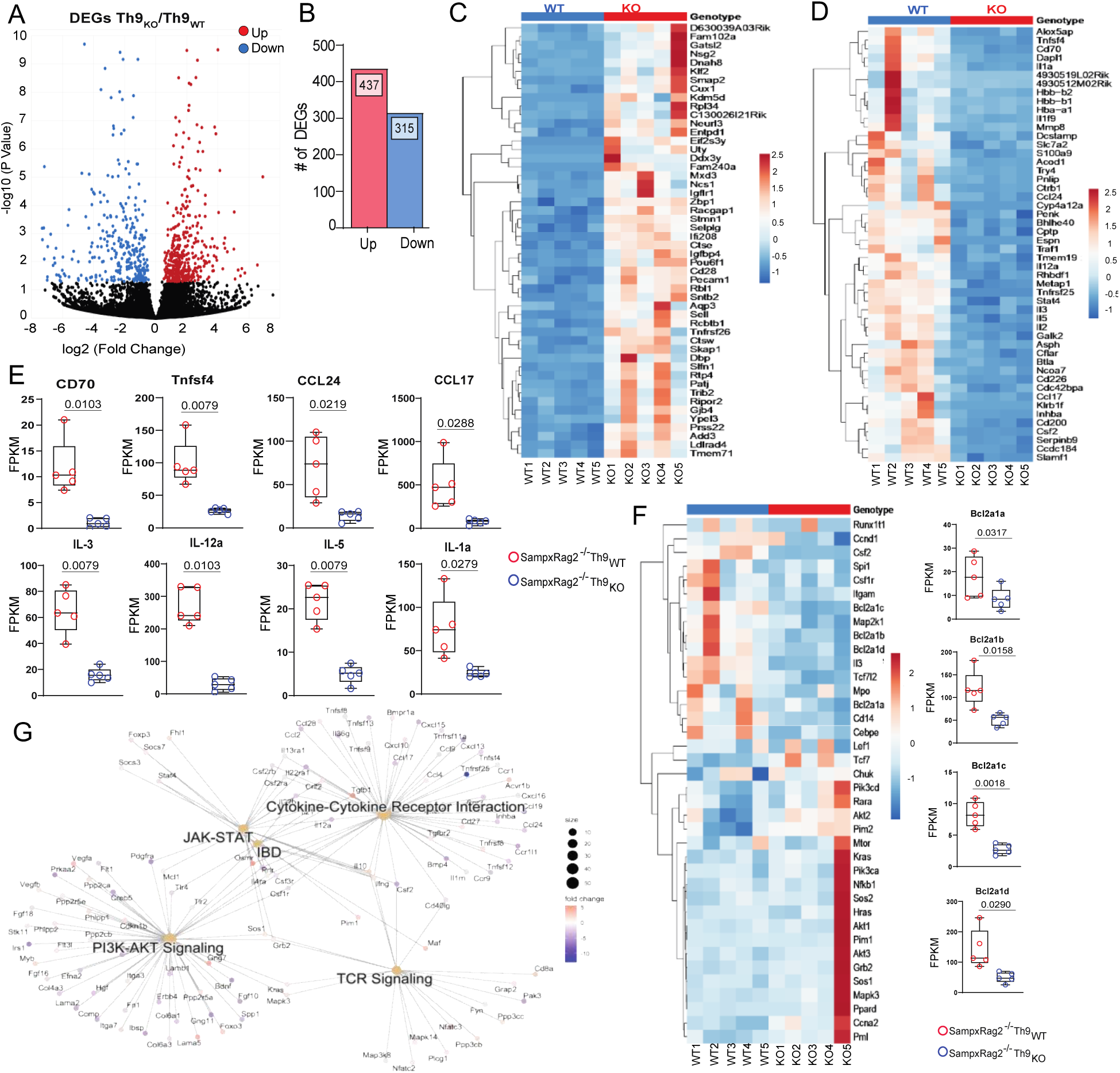
RNASeq analysis of TH9_WT_ and Th9_KO_ cells. **(A)** Volcano plot representation of DEGs in Th9_KO_ cells vs TH9_WT_ cells with fold difference between log2 normalized expression in Th9_KO_ vs TH9_WT_ cells plotted versus −log10 adjusted P-value. Red and blue points mark the genes with significantly increased or decreased expression respectively while black dots are gene with no significant difference in gene expression. (**B**) Bar chart displaying the number of significant DEGs up- and down-regulated in Th9_KO_ cells. (**C**) and (**D**) Heatmaps of top 50 differentially up- and down-regulated genes respectively, from RNA-seq of Th9_KO_ versus Th9_WT_ cells. Rows are centered for each gene with unit variance scaling applied, gene clustering by correlation distance and average linkage. (**E**) Comparative expression of gene of interest expressed as FPKM for each replicate. (**G**) Heatmap of the most significant DEGs in the IBD pathway. Rows are centered for each gene with unit variance scaling applied, gene clustering by correlation distance and average linkage. (**F**) Network analysis generated with Enrichplot and ClusterProfiler demonstrating relative relationship among expressed genes with log2(F) >1 and central nodes of IBD, TCR, JACK-STAT, PI3K-AKT signaling and Cytokine-Cytokine receptor signaling.

**Table 1:** KEEG pathways analysis. Table showing the top 20 significantly enriched KEGG pathways of differentially expressed genes (DEGs) between Th9_KO_ and Th9_WT_ cells an FDR value cutoff of 0.05.

| Pathway | DEGs | Genes in Pathway | FDR Pvalue |
| --- | --- | --- | --- |
| Cytokine-cytokine receptor interaction | 35 | 249 | 0.00038037 |
| Malaria | 13 | 54 | 0.00038037 |
| Cellular senescence | 23 | 170 | 0.00042099 |
| Pathways in cancer | 47 | 510 | 0.00051531 |
| Inflammatory bowel disease | 13 | 60 | 0.00051531 |
| Transcriptional misregulation in cancer | 21 | 182 | 0.00283391 |
| Pancreatic secretion | 15 | 104 | 0.00283391 |
| Apoptosis | 18 | 135 | 0.00348736 |
| African trypanosomiasis | 7 | 36 | 0.00377852 |
| Acute myeloid leukemia | 11 | 70 | 0.00412024 |
| Colorectal cancer | 12 | 86 | 0.00774856 |
| T cell receptor signaling pathway | 13 | 102 | 0.00774856 |
| Influenza A | 17 | 157 | 0.00778288 |
| Cell adhesion molecules | 18 | 154 | 0.00936057 |
| Human T-cell leukemia virus 1 infection | 23 | 231 | 0.00936057 |
| Cell cycle | 16 | 123 | 0.00936057 |
| Hepatocellular carcinoma | 17 | 165 | 0.00936057 |
| Asthma | 6 | 23 | 0.00945039 |
| AGE-RAGE signaling pathway in diabetic complications | 12 | 101 | 0.01132729 |
| Hepatitis B | 17 | 153 | 0.01498423 |
| Type I diabetes mellitus | 9 | 57 | 0.01498423 |
| JAK-STAT signaling pathway | 17 | 146 | 0.01498423 |
| Intestinal immune network for IgA production | 8 | 40 | 0.01498423 |
| Tuberculosis | 17 | 160 | 0.02268277 |
| C-type lectin receptor signaling pathway | 11 | 104 | 0.02564749 |
| Allograft rejection | 9 | 52 | 0.02786477 |
| Chemokine signaling pathway | 18 | 179 | 0.02786477 |
| Amoebiasis | 13 | 102 | 0.02945227 |
| Toxoplasmosis | 11 | 107 | 0.03100625 |
| Pancreatic cancer | 10 | 76 | 0.03247673 |
| Leishmaniasis | 10 | 69 | 0.03247673 |
| Rheumatoid arthritis | 11 | 83 | 0.03312427 |
| Measles | 14 | 132 | 0.03409195 |
| Autoimmune thyroid disease | 9 | 56 | 0.03750883 |
| Platinum drug resistance | 9 | 78 | 0.03919896 |
| NF-kappa B signaling pathway | 12 | 100 | 0.04045021 |
| MicroRNAs in cancer | 16 | 161 | 0.04213949 |
| FoxO signaling pathway | 14 | 126 | 0.0441537 |

To further explore which pathways are perturbed in Th9_KO_ cells, we used iPathwayGuide (Advaita Bioinformatics, Ann Arbor, MI) to perform Impact Analysis^26 27^. Among the most significantly perturbated pathways we found Inflammatory bowel disease (KEGG: 05321) that confirmed downregulation of IL-1α, IL-5 and IL-22 proinflammatory cytokines and upregulation of IL-10 and FOXP3 in Th9_KO_ cells (Fig. S 3A-B). In all, these findings are in line with our central hypothesis that DR3 deficient Th9 cells harbor a diminished inflammatory signature.

### Comparative phosphoproteomic analysis between Th9_WT_ and Th9_KO_ cells

To further explore DR3-driven signaling pathways in the Th9 cells network we performed a global analysis of protein phosphorylation in both Th9_WT_ and Th9_KO_ cells. More than 3000 sites of phosphorylation were identified, with 219 being differentially phosphorylated between Th9_WT_ and Th9_KO_ cells (Fig. 3A). The top 15 up- and down-expressed phosphoprotein in the Th9_WT_ vs the Th9_KO_ cells are depicted in Fig. 3C. We performed Ingenuity Pathway Analysis which revealed that Th9_WT_ cells displayed significantly activated pathways related to cytokine signaling, cellular cycle and proliferation and cellular immune response. Conversely, only one pathway (HIPPO signaling pathway, z score=-2.714) was found significantly inhibited when compared to the KO group (Fig 3B). Among the significant canonical pathways that were increased within the Th9_WT_ group, the IL-7 and IL-8 signaling pathways (z-score=2.121 and z-score=2.111 respectively) were of particular interest. Notably, IL-7 signaling determines chromatin modifications that ultimately increase accessibility of the *Il9* promoter locus and binding with the positive regulator FOXO1^28^.

**Figure 3:**
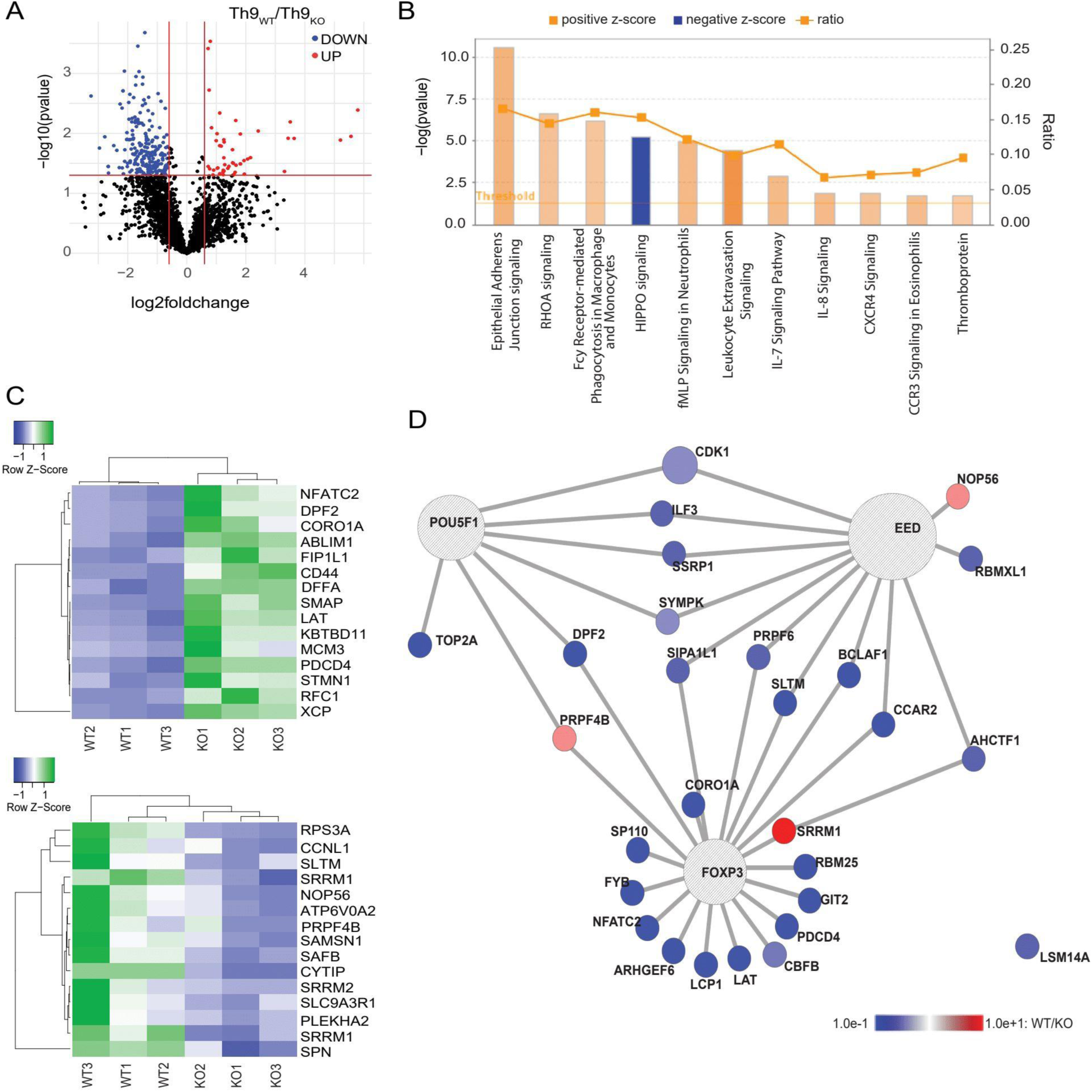
Phosphoproteomic analysis of TH9_WT_ and Th9_KO_ cells. **(A)** Volcano plot representation of differentially significant phosphopeptides in Th9_WT_ cells vs TH9_KO_ cells with fold difference between log2 normalized expression in Th9_WT_ vs TH9_KO_ cells plotted versus −log10 adjusted P-value. Red and blue points mark the genes with significantly increased or decreased expression respectively while black dots are gene with no significant difference in phosphorylation (**B**) Significant canonical pathways enriched in TH9_WT_ vs Th9_KO_ derived from IPA “Core Analysis”. Blue bars: negative z-score; orange bars: positive z-score, “Threshold” indicates the minimum significance level (scored as – log [P-value] from Fisher’s exact test, set here at 1.3). while “Ratio” (orange squares) indicates the number of molecules from the data set that map to the pathway listed divided by the total number of differentially expressed molecules found in each pathway over the total number of proteins in that pathway. (**C**) Heat map of top 15 up- and down-expressed phosphoprotein in Th9_WT_ vs Th9_KO_ cells. (**D**) Network plot indicating experimentally measured and predicted intra-protein crosstalk of selected phosphopeptides that are significant in our data set. Crosstalker designates each node with a distinct color on the basis of rate ratio between Th9_WT_ vs Th9_KO_ cells. The blue and red colors indicate phosphopetide downregulation or upregulation respectively (n=3).

Overall, these results support our hypothesis that Th9 cells with functional DR3 have a proinflammatory signature in our mouse model. Along with our RNA-seq data, the phosphoproteome analysis revealed major differences in the expression of phosphopetides involved in the TCR signaling. For instance, significant dephosphorylation of NFACT2, LAT, CORO1A was present in Th9_WT_ compared to Th9_KO_ cells. Activation of this proteins following TCR ligation is required for optimal T cell development, differentiation and activation^29^, while defects in TCR signaling molecules have been linked to varying degrees of autoimmune conditions^30^.

An additional approach to examining the correlation between the most significant phosphopetides in our data set was performed using Crosstalker (YourOmics, Inc.; www.youromics.com), a novel software analysis tool^31^, which helps to reveal previously unidentified targets that may play a key role in maintaining the phenotype. Our analysis showed that, overall, proteins that are linked to the FOXP3 network were functional in Th9_KO_ cells but frequently inactivated in Th9_WT_ cells (Fig. 3D). This result is in line with our previous work that showed DR3 stimulation targets FoxP3^+^ cells and converts regulatory cells to dysfunctional CD25^−^FoxP3^+^cells, which accelerates disease manifestation in our SAMP mice^11^.

### DR3-deficient Th9 cells have diminished *in vivo* colitogenic potential in adoptive transfer models of intestinal inflammation

In our next step, we sought to investigate the function of Th9-polarized cells *in vivo.* We utilized two adoptive transfer models of intestinal inflammation to test our hypothesis that Th9 cells that are deficient in DR3 have diminished ability to induce intestinal inflammation to recipient mice. First, we exploited the SAMP×Rag2^-/-^ adoptive transfer model to compare the pathogenic potential of donor, SAMP-derived Th9 cells that had either intact (Th9_WT_) or deficient (Th9_KO_) DR3 signaling (Fig. 4A). We observed that mice reconstituted with Th9_WT_ cells had significantly more severe colitis and higher (although not statistically significant) inflammatory scores at the ileum than recipients of Th9_KO_ cells (Fig. 4B, 4C). Furthermore, we observed significantly lower gene expression of pro-inflammatory cytokines IL-9, IL-1β and TNF in both colonic and ileal tissues from SAMP×Rag^-/-^ reconstituted with Th9_KO_ compared to recipients from Th9_WT_ cells (Fig 4D, 4E). In contrast, IL-6 was markedly increased only in the colon of mice recipient of Th9_KO_ cells (Fig 4D), whereas, in both locations, we detected increased levels of IL-10 in mice reconstituted with polarized-Th9 lacking functional DR3 (Fig 4D, 4E). Alongside those results, we also observed the same expression patterns in both colon and ileum regarding genes involved in Th9 differentiation such as Baft3 and Id3, which were down and up-regulated respectively (Fig 4D, 4E). Finally, FACS analysis of CD4-gated lymphocytes isolated from spleens of SAMP×Rag2^-/-^ mice showed higher frequency of CD3+CD4+ T cells producing IL-9 in mice reconstituted with Th9 _WT_, while splenocytes from mice reconstituted with Th9_KO_ cells displayed a marked increase of IL-10 producing cells (Fig. 4F).

**Figure 4:**
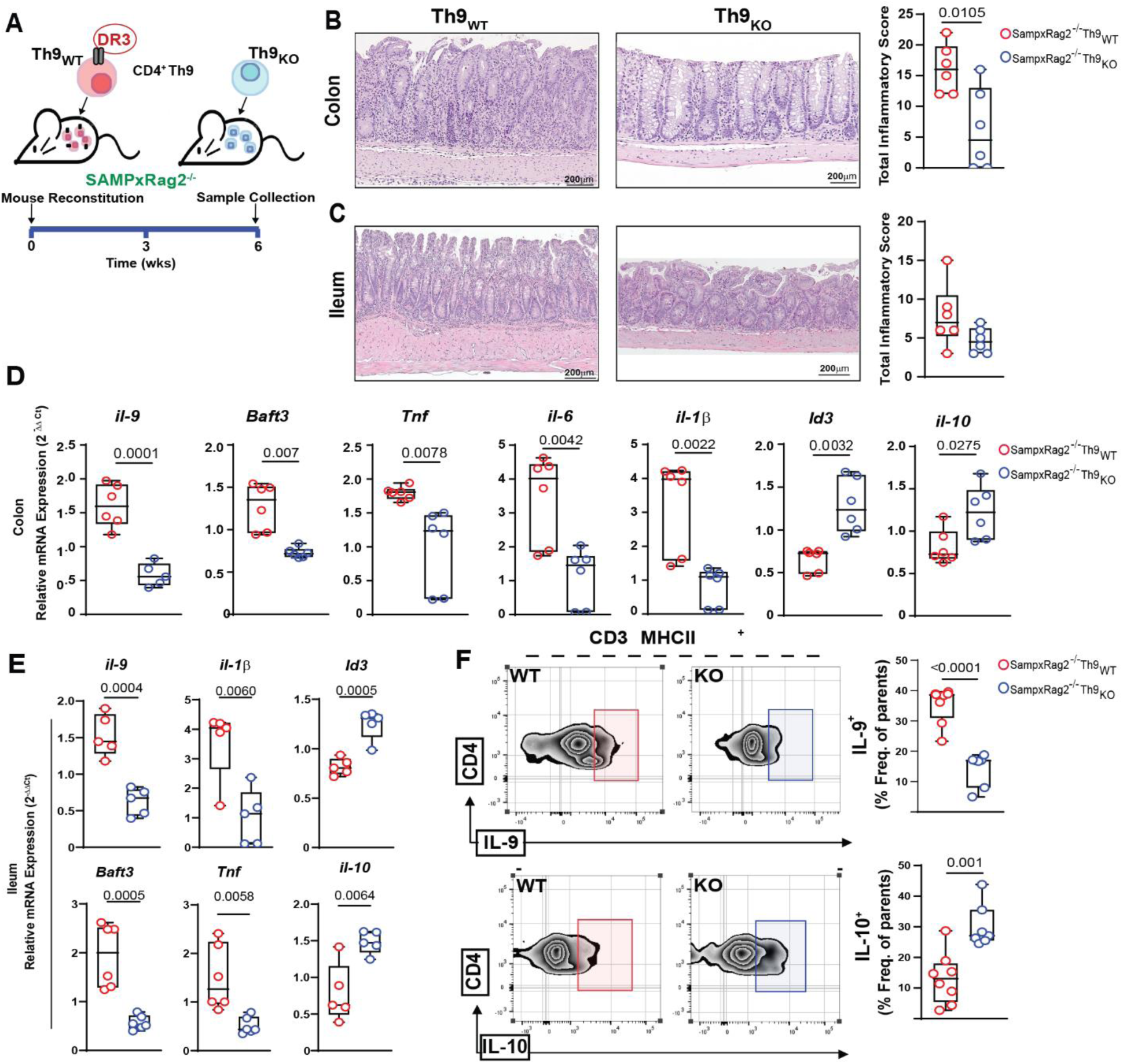
Adoptive transfer of Th9 with functional DR3 induces colitis in SAMPxRag^-/-^ recipients and leads to worse ileitis. (**A**) Experimental workflow adoptive transfer model using Th9_WT_ and TH9_KO_ (**B**) and (**C**) Representative images of full-thickness H&E-stained colon and ileal tissues, scale bar= 200µm with histologic analysis (reported as total inflammatory score) of colon and ileum respectively from SAMPxRag2^-/-^ adoptively transferred with Th9_WT_ or TH9_KO_. (**D**) and (**E**) Relative expression of indicated gene mRNAs measured in total tissue from colon and ileum of WT and KO mice. The mRNA levels were determined by RT-qPCR, normalized to β-actin and expressed as fold change (2−^ΔΔCt^). (**F**) Representative FACS histograms showing intracellular IL-9 and IL-10 expression among CD4-gated lymphocytes isolated from spleens of Rag2^-/-^ mice adoptively transferred with Th9_WT_ or TH9_KO_. Data, indicated as mean±SD, correspond to 2 independent experiments; statistical analysis was determined by, 2-tailed Unpaired t-test, n= 6).

To further substantiate our findings, we repeated our adoptive transfer experiments, using Rag2^-/-^ mice as recipients (Fig. 5A). In line with our previous results above, our study clearly showed that Th9_KO_ cells were less pathogenic than Th9_KO_ as they induced significantly less severe colitis to recipient mice (Fig. 5B-D). We also observed similar gene expression patterns, since colonic tissue from recipients of Th9_KO_ cells exhibited decreased mRNA levels of pro-inflammatory cytokines such as IL-6, TNF-α, INF-γ, IL-1β and il-9 as well, and higher IL-10 mRNA expression (Fig. 5E). We also observed a striking increase in expression of genes associated with negative regulation of IL-9 such as STAT6, Tak1 and Id3 in colon tissue from mice reconstituted with Th9_KO_ cells. Conversely, Baft3, which has been proven to be an inductor of Th9 cell differentiation was markedly decreased in Rag2^-/-^ recipients of Th9_KO_ (Fig. 5E). Finally, mice reconstituted with Th9_WT_ cells mice harbored spleens enriched in higher frequency of CD3^+^CD4^+^ T cells producing IL-9, while splenocytes from mice reconstituted with Th9_KO_ cells displayed a marked increase of IL-10 producing cells (Fig. 5F).

**Figure 5:**
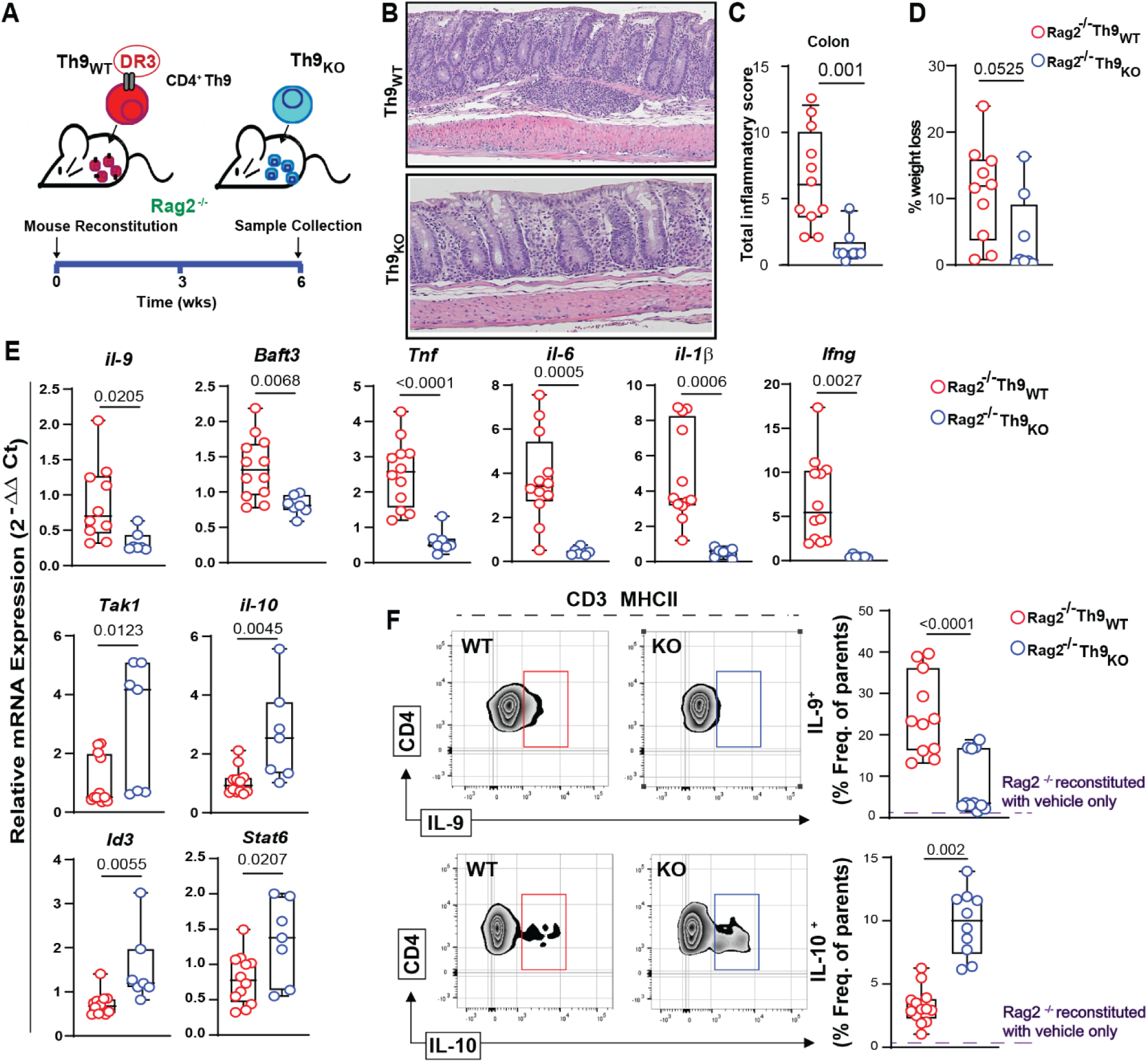
Adoptive transfer of Th9 from DR3-competent—but not DR3-deficient— SAMP mice induces significant colitis in Rag2^-/-^ recipients. (**A**) Experimental workflow adoptive transfer model using Th9_WT_ and TH9_KO_ (**B**) Representative images of full-thickness H&E-stained colon tissues, scale bar= 200µm (**C**) Histologic analysis (reported as total inflammatory score) of colon tissue from Rag2^-/-^ adoptively transferred with Th9_WT_ orTH9_KO_ (**D**) Percentage of total body weight loss in Rag2^-/-^ mice recipients from Th9_WT_ cells compared to in Rag2^-/-^ mice recipents from Th9_KO_ (**E**) Relative expression of indicated gene mRNAs measured in total tissue from the ileum of WT and KO mice. The mRNA levels were determined by RT-qPCR, normalized to β-actin and expressed as fold change (2−^ΔΔCt^). (**F**) Representative FACS histograms showing intracellular IL-9 and IL-10 expression among CD4-gated lymphocytes isolated from spleens of Rag2^-/-^ mice adoptively transferred with Th9_WT_ or TH9_KO_. Data, indicated as mean ± SD, correspond to 2 independent experiments; statistical analysis was determined by, 2-tailed Unpaired t-test, n≥ 7).

Taken together, these results show that DR3 deficiency critically affects the function of Th9 cells in SAMP mice by diminishing their negative regulation and ability to produce IL-9. This defect is associated with a reduced pathogenic potential of these cells and protection from inflammation when adoptively transferred.

### Blockade of IL-9 ameliorates CD-like ileitis in SAMP mice

Based on our findings that showed decreased pathogenic potential of DR3-deficient Th9 cells and literature reports of anti-inflammatory effects of anti-IL-9 blockade in experimental models of colitis^32, 33^, in our final experiment we tested whether neutralization of IL-9 would ameliorate inflammation in SAMP1/YitFc mice. This could be of clinical importance as, differently from most other models, SAMP mice develop terminal ileitis spontaneously that shares several similarities with CD; thus our results could be transferable to the human condition. To investigate whether IL-9 blockade could effectively ameliorate intestinal inflammation in ileitis-prone SAMP mice, we inhibited IL-9 function in gender-matched 20-week-old SAMP mice by intraperitoneal injection of 100μg anti-IL-9, or an isotype-matched control Ab (IgG), twice a week for 4 weeks. Our results clearly showed that IL-9 blockade is an effective treatment for CD-like terminal ileitis in SAMP mice. Indeed, mice that were administered the neutralizing anti-IL-9 mAb demonstrated a significant reduction of total inflammatory score in the ileum, as compared to isotype-treated controls (Fig. 6A). This clinicopathological effect was coupled with downregulation of pro-inflammatory cytokines gene expression after treatment of SAMP mice with anti-IL-9 (with the exception of IL-6) (Fig. 6B). Taken together, these results provide mechanistic evidence for the potential of anti-IL-9 administration as an effective anti-inflammatory treatment in patients with CD.

**Figure 6:**
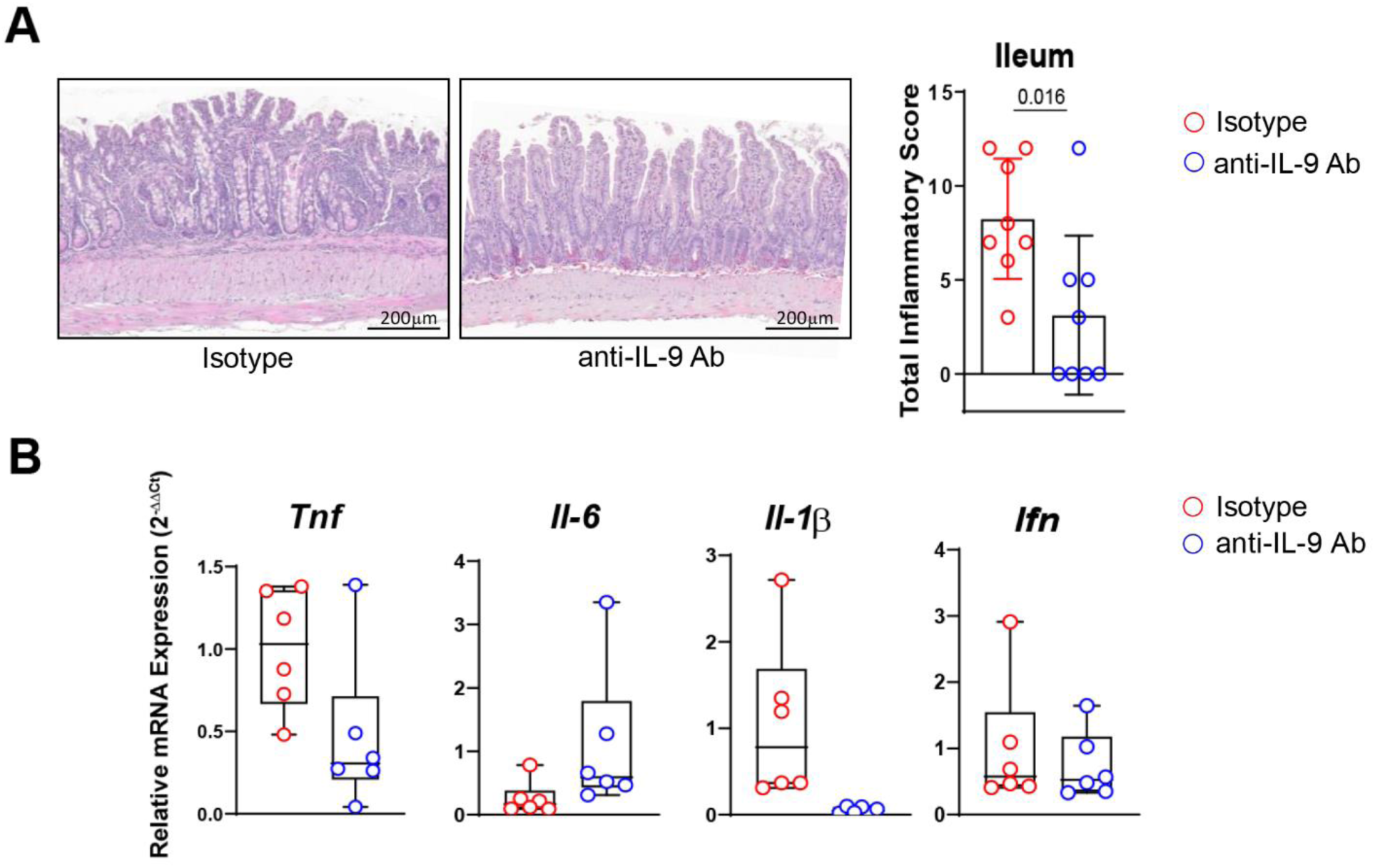
Blockade of IL-9 ameliorates CD-like ileitis in SAMP mice. (**A**) Representative images of full-thickness H&E-stained ileal tissues, scale bar= 200µm with histologic analysis (reported as total inflammatory score) of ileum from SAMP mice treated with aIL-9 ab compared to SAMP mice treated with Isotype control. (**B**) Relative expression of indicated gene mRNAs measured in total tissue from the ileum of treated vs control mice. The mRNA levels were determined by RT-qPCR, normalized to β-actin and expressed as fold change (2−^ΔΔCt^).

## Discussion

In the present study, we provide evidence for a pivotal role of TL1A/DR3 signaling in the function of Th9 cells with implications for the pathogenic potential of the latter during chronic intestinal inflammation. We show that TL1A/DR3 strengthen the polarization of and IL-9 production by Th9 lymphocytes and demonstrate that signaling through DR3 confers a pro-inflammatory phenotype to Th9 cells, with high production of relevant cytokines. Furthermore, by transcriptomic and phosphoproteomic analyses, we show that stimulation via TL1A induces distinct inflammatory signatures in Th9 cells, including several that are associated with immunological pathways that are critical for IBD. Finally, we provide mechanistic data for the existence of an immunological TL1A/DR3/Th9 axis that mediates chronic intestinal inflammation in animal models of CD, by showing decreased capability of DR3-deficient Th9 cells to adoptively transfer ileocolitis and anti-inflammatory effects of IL-9 neutralization.

Our findings clearly support a major role of DR3 signaling in both the differentiation of Th9 cells and their pathogenic signature in chronic CD-like intestinal inflammation. To test this dual hypothesis, we utilized the SAMP1/YitFc mouse as the appropriate research tool, since, on the one hand, it develops spontaneous ileitis with remarkable pathological and immunological similarity to human CD^17, 34^ and, on the other hand, both TL1A and DR3 are highly expressed in SAMP mice during the chronic phase of the disease ^12, 14^. This upregulation bears pathogenetic significance, as we have previously shown (and confirmed herein) that SAMP mice with a deletion of the *dr3* gene (DR3^-/-^xSAMP) are protected from severe ileitis^15^. The study of DR3^-/-^xSAMP mice allowed us to identify a significant role of DR3 in the differentiation of Th9 cells. First, deletion of *dr3* is associated with reduced expression of IL-9 at the terminal ileum both at the mRNA and protein levels. Second, in the absence of DR3, a rearrangement takes place regarding the expression of Th9-related regulatory factors, with diminished expression of Th9 inducers and upregulation of Th9 inhibitors, such as *Gata3*, *Stat6*, *Tak1*, *Id3*. Finally, in our *in vitro* model we clearly showed that the addition of TL1A to a Th9 polarizing microenvironment significantly expands the Th9 pool and increases the secreted IL-9 protein. In fact, in our system the addition of TL1A in the cell cultures appeared to be critical for the acquisition of the Th9 phenotype by lymphocytes. Consistent with our findings, the TL1A/DR3 system is also involved in promotion of IL-9-producing T (Th9) cells differentiation in a model of allergic lung inflammation^35^, with enhanced IL-9 production. Similarly, Meylan et al. provided evidence for a regulatory role of DR3 in Th9 cell function and promotion of IL-9 secretion in a model of allergic disease showing that augmentation of secreted IL-9 depends on DR3 signaling^36^.

Our present study indicates that, in addition to augmenting Th9 generation and IL-9 secretion, TL1A/DR3 signaling appears to significantly augment the pro-inflammatory potential of this lymphocytic population. In separate lines of evidence, it has been shown that both TL1A/DR3 and TH9/IL-9 confer pathogenicity in clinical and experimental chronic intestinal inflammation. On the one hand, it is well established that, upon TL1A binding, DR3 promotes T cell accumulation and pathology in inflammatory disease models^36^. In addition, the TL1A/DR3 system is highly upregulated in involved areas of patients with IBD and bear functional significance^12, 37^. On the other hand, emerging evidence also suggests a functional role of Th9 cells in IBD^38^ through secretion of IL-9 which is responsible of impairing intestinal barrier integrity and repair^39, 40^. Herein, we provide data which supports the linking of the two pathways via the stimulatory effect of activated DR3 on Th9 pathogenicity, as shown by the distinct phenotypes between SAMP-derived Th9 cells that were either functional (Th9_WT_) or deficient (Th9_KO_) in DR3 signaling, in particular, a prominent pro-inflammatory signature of the Th9_KO_ population. This is supported by several findings presented herein. Firstly, Th9_WT_ cells displayed not only significantly increased secretion of IL-9, but of other pro-inflammatory cytokines, as well, including the IBD-definitive TNF. Conversely, Th9_KO_ showed increased production of the anti-inflammatory cytokine IL-10. Secondly, our transcriptomic analysis confirmed that pro-inflammatory pathways were significantly depleted in Th9_KO_ as compared to Th9_WT_ cells. Of great importance, the IBD pathway was one of the most differentially expressed between the 2 groups. Particularly, DR3-deficient Th9 cells displayed increased levels of the anti-inflammatory cytokine IL-10 and FOXP3 transcription factor which is a master regulator of the development and functions of Treg cells. We also show upregulation of TGFβ1, a pleiotropic cytokine with potent regulatory activity which has been demonstrated to induce Foxp3 expression in CD4^+^CD25^−^ naive T cells to enforce transition to regulatory T cells^41^. Conversely, within the same pathway, pro-inflammatory cytokines (IL-12, IL-1α, IL-22 and IL-5) are downregulated in Th9_KO_ versus Th9_WT_ cells. Finally, multiple genes associated to inflammatory processes and IBD are upregulated in Th9_WT_ (*BCL2, CFS2* and *CSF1r*). Thirdly, our phosphoproteomic analyses, which, to our knowledge is the first to be carried out on Th9 cells, uncovered a significant activation of pathways related to cytokine signaling, and cell migration in Th9_WT_ cells compared to DR3-deficient Th9 cells. Specifically, the activation of IL-8 and CCR3 pathways play an important role in the expression of adhesion molecules (*eg.* Integrins) and T cells trafficking^42^. Multiple lines of evidence have shown that IL-8 is a cytokine that is significantly overexpressed in affected intestinal areas from UC and CD patients^43, 44^ and it is involved in the pathogenesis of UC^45^. Similarly, the CC chemokine receptor 3 (CCR3) may be of pathogenetic importance in IBD, as specific blockade of CCR3 in the spontaneous eosinophilic CD-like SAMP1/SkuSlc mouse with a monoclonal antibody resulted in a striking reduction of ileitis^46^. Interestingly, anti-trafficking therapeutic approaches have been approved for the treatment of patients with IBD^47^. Finally, our adoptive transfer experiments present definitive mechanistic proof of the pathogenic potential of pro-inflammatory, TL1A/DR3-induced Th9 cells, since cells that were deficient in DR3 signaling were also deficient in inducing severe ileo-colitis in recipient RAG2-/- mice.

Overall, our results show that DR3 stimulation sets the Th9 immunostat towards the pro-inflammatory mode. Converging lines of evidence indicate that this novel inflammatory pathway may be of pathogenetic importance and offer new therapeutic options for patients with CD. Recent data points to a positive association between IL-9 levels and disease activity^48^ in CD. Importantly, we also report herein a highly significant correlation between IL-9 and ileitis severity in SAMP mice, a finding that further emphasizes the similarity between the human condition and this mouse model. Moreover, neutralization of IL-9 via the administration of a monoclonal antibody resulted in amelioration of disease severity in SAMP mice, further supporting the biological importance of this pathway for the development of IBD-reminiscent intestinal inflammation. Given the fact that the SAMP is considered one of the few experimental models of ileitis/CD, our results lead to the appealing idea of targeting this pathway as a treatment option for the human condition. This is further supported by the strong anti-inflammatory effect of anti-IL-9 treatment in SAMP mice with ileitis. Therefore, our results raise the possibility that a similar therapeutic approach in patients with IBD may prove beneficial. Supporting this hypothesis is the fact that anti-TL1A therapeutic strategies are currently underway in IBD with satisfactory results been presented so far^16^.

Overall, our results agree with previous studies from our laboratory which identified DR3 signaling as a modulator of a multicellular network, including Tregs, in ileitis-prone SAMP mice. More importantly, it reveals a novel role of DR3 in Th9 cells development that appears to regulate their pro-inflammatory phenotype in models of CD-like ileitis and colitis. This finding is strongly supported by our adoptive transfer studies where we observed that Th9_WT_ cells were able to induce a more severe intestinal inflammation in both Rag2^-/-^ and SAMP×Rag2^-/-^ compared to the Th9_KO_ cells and by the beneficial effects of IL-9 blockade in SAMP CD-ileitis. These findings indicate that TLA1/DR3 and Th9 cells may be valid therapeutic targets in CD.

## Material and Methods

### Animals

AKR, SAMP and DR3-/-×SAMP and SAMP×Rag2-/- colonies are maintained under SPF conditions at the CWRU Animal Facility. Mice are routinely tested for common pathogens. Rag2-/- [C.129S6(B6)-Rag2tm1Fwa N12] mice are purchased from Jackson Laboratories (Bar Harbor, Maine). Animals have access to food ad libitum and are maintained on a 12-hour light/dark cycle.

### Histology

Murine ileum and colon samples were collected, rinsed with PBS, opened longitudinally, formalin-fixed (Fisher Scientific, Pittsburgh, PA), and paraffin-embedded. Subsequently tissues were sectioned at 3–4μm and stained with hematoxylin and eosin (H&E). Histological evaluation of inflammation severity was determined by using a semiquantitative scoring system as previously described^49^.

### Stereomicroscopy

Ileal tissue abnormalities (i.e., cobblestone lesions) and normal mucosa were investigated by examining the cellular structural pattern ileal tissue via stereomicroscopy, cm by cm, using a reference catalogue of lesions, as previously described^50^. Both healthy and cobblestone-like areas were calculated per cm using ImageJ software (NIH, Bethesda, MD, USA).

### Isolation of splenocytes (SPLs)

SPLs were isolated as described previously^11^. The resulting cell suspension was depleted of erythrocytes and used for downstream applications.

### Isolation and culture of CD4+ T cells

Spleens were removed aseptically and gently pressed against a 100-µm cell strainer to obtain single-cell suspensions. Enriched CD4+ cells were obtained by magnetic sorting, using a CD4^+^ T-cell isolation kit from Miltenyi Biotec Inc. (Auburn, CA). Purified CD4^+^ cells were cultured in 24-well round-bottom plates (1×106 cells/well) with plate-bound anti-CD3e (2.5 μg/ml, clone 2C11, BD Biosciences, San Diego, CA), soluble anti-CD28 (2 µg/ml, clone 37.51, BD Biosciences, San Diego, CA), recombinant IL-4 (20 ng/ml, R&D) and TGF-β (3 ng/ml, R&D) in RPMI 1640 medium with 10% FBS, 2 mM L-glutamine, 10 mM β-mercaptoethanol, and 1% penicillin/streptomycin. Recombinant IL-2 (20 ng/ml, R&D) was added after 24h to cell cultures. Cells and supernatants were collected after incubation for 72h at 37°C and 5% CO_2_ and stored at −80°C until further use.

### Generation of Th9 cells

CD4+ cells were enriched from SPLs by negative selection using CD4 microbeads on an AutoMACs separator, according to manufacturer’s instructions (Miltenyi Biotech, Cambridge, MA). Naïve T cells were then sorted by FACS using combination of markers (-CD3, -CD4, -CD44, -CD62, Zombie viability dye). Purified CD4^+^ T cells were cultured in 6-well round-bottom plates (1×10^6^ cells/well) with RPMI complete media supplemented with plate-bound anti-CD3ε (2.5 μg/ml; Clone 145-2C11; eBioscience) and soluble anti-CD28 (2.5 μg/ml; Clone 37.51; eBioscience) for 3 days under polarizing conditions for Th9 (20 ng/ml rIL-4, 4 ng/ml rTGF-β,) with or without addition of 10 ng/ml TL1A). Cells with medium only were used as control (Th0). Cell number, phenotype and purity were confirmed by flow cytometer analysis.

### Cytokine measurements

The concentrations of multiple cytokines in the supernatants from tissue and cell cultures were measured by ELISA (eBioscience) or Q-Plex Array Mouse cytokine screen IR Quansys16-Plex kit (Quansys Biosciences, Logan, UT), according to the manufacturer’s instructions.

### Real-time RT-PCR

Total RNA was extracted from homogenized ileal and colonic tissues by use of the RNeasy Mini kit (Qiagen, Valencia, CA) and reverse transcribed (High-Capacity cDNA Reverse Transcription Kit; Applied Biosystems, Forest City, CA), both according to the manufacturer’s instructions. Quantification of gene expression was performed on Light-Cycler® 480 System using SYBR Green methodology, according to the manufacturer’s instructions. Relative mRNA expression of each target gene was normalized β-actin and calculated by the ΔΔCt method.

### In vivo treatment of mice with anti-IL-9

Mice were injected with 100 μg anti-IL-9 (MM9C1; BioXCell) intraperitoneally twice a week for 4 weeks. Control mice were injected with same amount of isotype control.

### Adoptive Cell Transfer

Differentiated Th9 cells (0.5×10^5^) from naïve CD4^+^ T cells of spleens of 20-wks SAMP and SAMPxDR3^-/-^ mice were collected and adoptively transferred by i.p. injection into 12-weeks-old Rag2^-/-^ and SAMP×Rag2^-/-^ mice as previously described. Recipient mice were euthanized 6 weeks after the transfer. Body weight loss and general malaise were monitored till euthanasia. Colon and ileum were excised for histological assessment of inflammation. Intestinal tissues were collected for gene and protein expression profiling.

### RNA-seq and data analysis

Total RNA was isolated and used for RNA-seq analysis, and the cDNA library was constructed by Beijing Genomics Institute using the Illumina HiSeq X platform (Shenzhen, China). High-quality reads were aligned to reference genome using Bowtie2^51^ and then gene expression level was calculated with RSEM^52^. For RNA-seq analysis, sequencing reads generated from the Illumina platform were assessed for quality and trimmed for adapter sequences using TrimGalore! v0.4.2 (Babraham Bioinformatics), a wrapper script for FastQC and cutadapt. Reads that passed quality control were then aligned to the human reference genome (GRCh38) using the STAR aligner v2.5.1. The alignment for the sequences were guided using the GENCODE annotation for GRCh38. The aligned reads were analyzed for differential expression using Cufflinks v2.2.1, a RNASeq analysis package which reports the fragments per kilobase of exon per million fragments mapped (FPKM) for each gene. Differential analysis report was generated using Cuffdiff. Differential genes were identified using a significance cutoff of q-value < 0.05 (FDR). The genes were then subjected to gene set enrichment analysis with iPathwayGuide (AdvaitaBio) for pathway analysis. This method uses two types of evidence: i) the over-representation of DEGs in a given pathway and ii) the perturbation of that pathway computed by propagating the measured expression changes across the pathway topology. The underlying pathway topologies, comprised of genes and their directional interactions, are obtained from the KEGG database^53, 54, 55^. Additional custom gene set enrichment analysis and visualization were performed in R using the DOSE, enrichplot, and fgsea packages^56, 57^.

### Unbiased Label Free Shotgun LC-MS/MS and data analysis

For this experiment a label-free 4-hour gradient LC-MS/MS protocol via an UPLC/TLQ-VELOS mass spectrometer was utilized, an approach that typically detects >800-1500 proteins per sample. Automated differential quantification of peptides was accomplished with Rosetta Elucidator, which is a software suite that automates qualitative and quantitative comparison of MS data sets, including algorithms for charge state determination, monoisotopic peak assignment, chromatographic peak construction, chromatographic alignment, and cross-experiment integration. Elucidator analysis involves deconvolution of multiple mass spectra taken across the chromatographic gradient. Multiple isotopic peaks, charge states and MS scans can be combined to extract the intensities of mass-chromatographic peaks in each LC-MS run, and converted into a single intensity value, corresponding to a distinct peptide species. The peptide peaks from multiple LC-MS runs are matched based on their mass/charge (m/z) ratio and elution times. Peptide and protein identifications are integrated with these peptide quantifications yielding a peptide expression matrix that can be used for subsequent statistical analysis. Protein and peptide teller algorithms with independent decoy validation are used for protein identifications and error analysis, and identifications are accepted if probability values are than ≤ 0.01 (equivalent to FDR). Data analysis was performed using Ingenuity Pathway Analysis (IPA; www.ingenuity.com, Qiagen, CA, USA) with the Core Analysis module to identify significant canonical pathways affected between Th9_WT_ and Th9_KO_ cells. Briefly, the regulated phosphoproteins, and their log2-transformed SILAC ratios between WT and KO samples were uploaded into the IPA software. Using the core analysis module, IPA generated computational networks that are biologically relevant to a priori defined functions. The mouse genes/protein names were converted to their human gene ortholog names for canonical pathway analysis. Pathways were considered significant based on the Fisher exact test with a –log (p-value) >1.3 (corresponds to a p-value <0.05), and a z-score >2 or <−2 was defined as the threshold of significant activation or inhibition, respectively. Additionally, we used Crosstalker (YourOmics, Inc.; www.youromics.com) software analysis tool^31^. In this computational approach our proteomic targets with significant fold change between WT and KO are used to “seed” a search for small protein-protein interaction sub-networks that are functionally associated with these targets. The identification of candidate sub-networks with a significant association to our targets can reveal proteins that were not originally identified as differentially expressed, but still exhibit significant crosstalk to our targets, which helps to reveal previously unidentified targets that may play a key role in maintaining the phenotype. To seed the analysis, only phosphopeptides with a p value <0.05, and a ratio change (WT/KO) of <0.5 or >2.0 were considered.

### Statistical analysis

Data were analyzed using GraphPad Prism 8, Phyton, R, Flowjo. Selection of appropriate statistical tests was based on variance and underlying distribution of data. Global effects between groups were assessed using one-way ANOVA with Bonferroni correction for multiple comparisons. Differences between individual groups were directly compared using two-sample unpaired Student’s t-test and results expressed as mean±SD, unless otherwise indicated, with P<0.05 considered significant.

## Acknowledgments

This work was supported by National Institutes of Health Grants DK042191, DK055812, and DK091222 awarded to Fabio Cominelli. We also acknowledge the Mouse Models Core and the Histology/Imaging Core of the Cleveland Digestive Disease Research Core Center (DK097948). We also thank Natalia Aladyshkina and Ashtyn Balasko for their management of the mouse colonies and acknowledge Dr. Daniela Schlatzer for her assistance with the phosphoproteome analysis. Finally, we would like to thank the Case Western Reserve University Genomics Core Facilities and Cytometry Core Facilities for their assistance with 16s rRNA sequencing and flow cytometry, respectively.

## Author Contributions

PM and LFB shared the first authorship and contributed to the acquisition, analysis, and interpretation of data, drafting of manuscript, and statistical analysis; FC contributed to the study concept and design, analysis and interpretation of data, drafting and critical revision of manuscript for important intellectual content, attainment of funding, and study supervision. AG contributed to the study design and to the drafting and critical revision of manuscript for important intellectual content. PM, LB, NA, AG, and K-AB performed the experiments. E.R.C. was responsible for computational analyses. TTP and GB contributed to the drafting and critical revision of manuscript for important intellectual content. All authors contributed to refinement of the study protocol and approved the final manuscript.

---

**Figure Suppl. 1:**
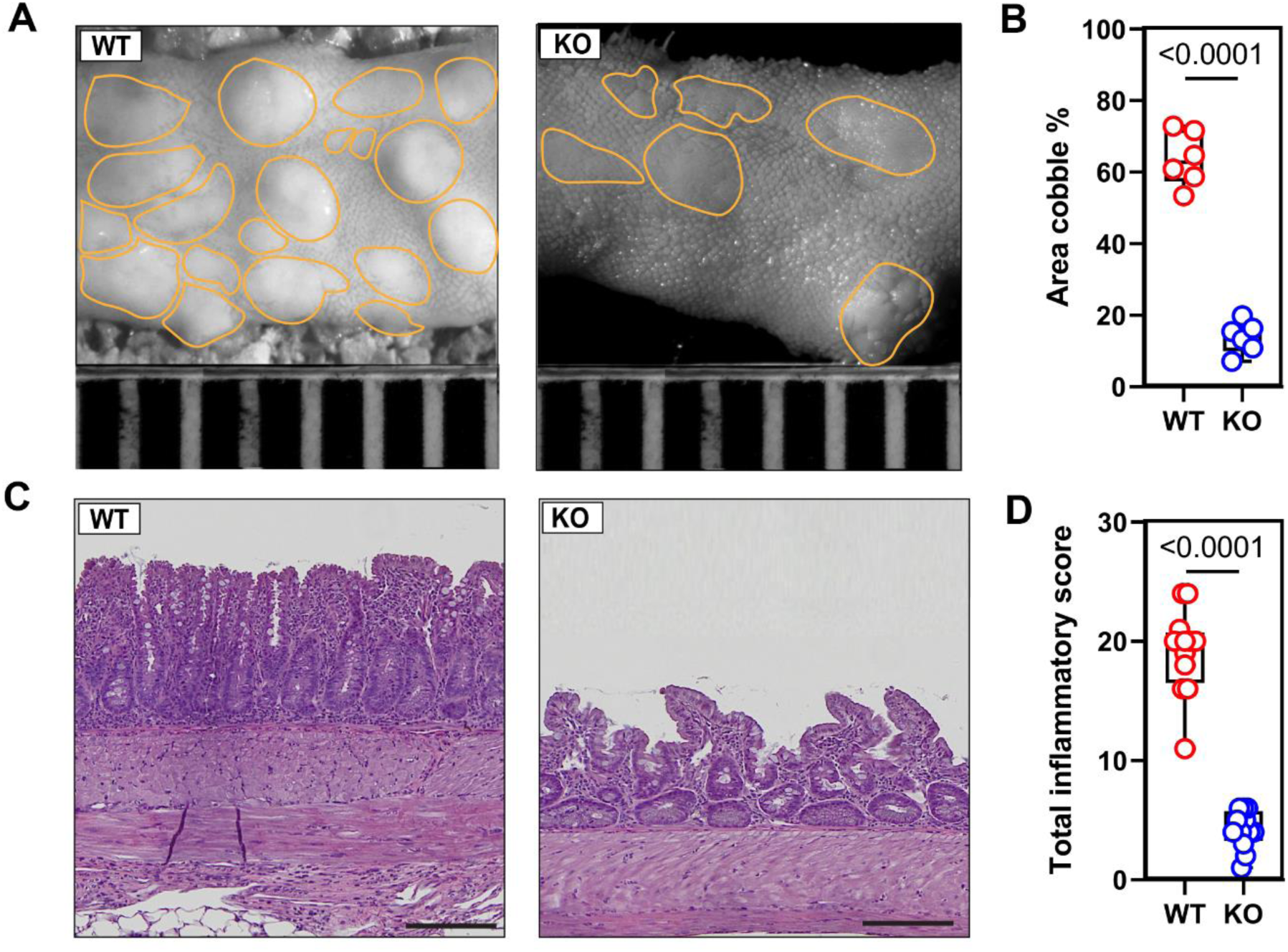
DR3^-/-^xSAMP (KO) mice exhibit a reduced inflammation compared to SAMP mice (WT). (**A**) Mucosal architecture of fixed postmortem ileal specimens collected from SAMP (WT) and DR3^-/-^xSAMP (KO) mice; (**B**) Cobblestone area expressed as percentage of total specimen calculated in the ileum of WT and KO mice (n=6). (**C**) Representative photomicrographs of ileal sections of 20-wk AKR and SAMP mice. Scale bar is 200 μm. (**D**) Total inflammatory score in 20-wks WT and KO mice (n=10).

**Figure Suppl. 2:**
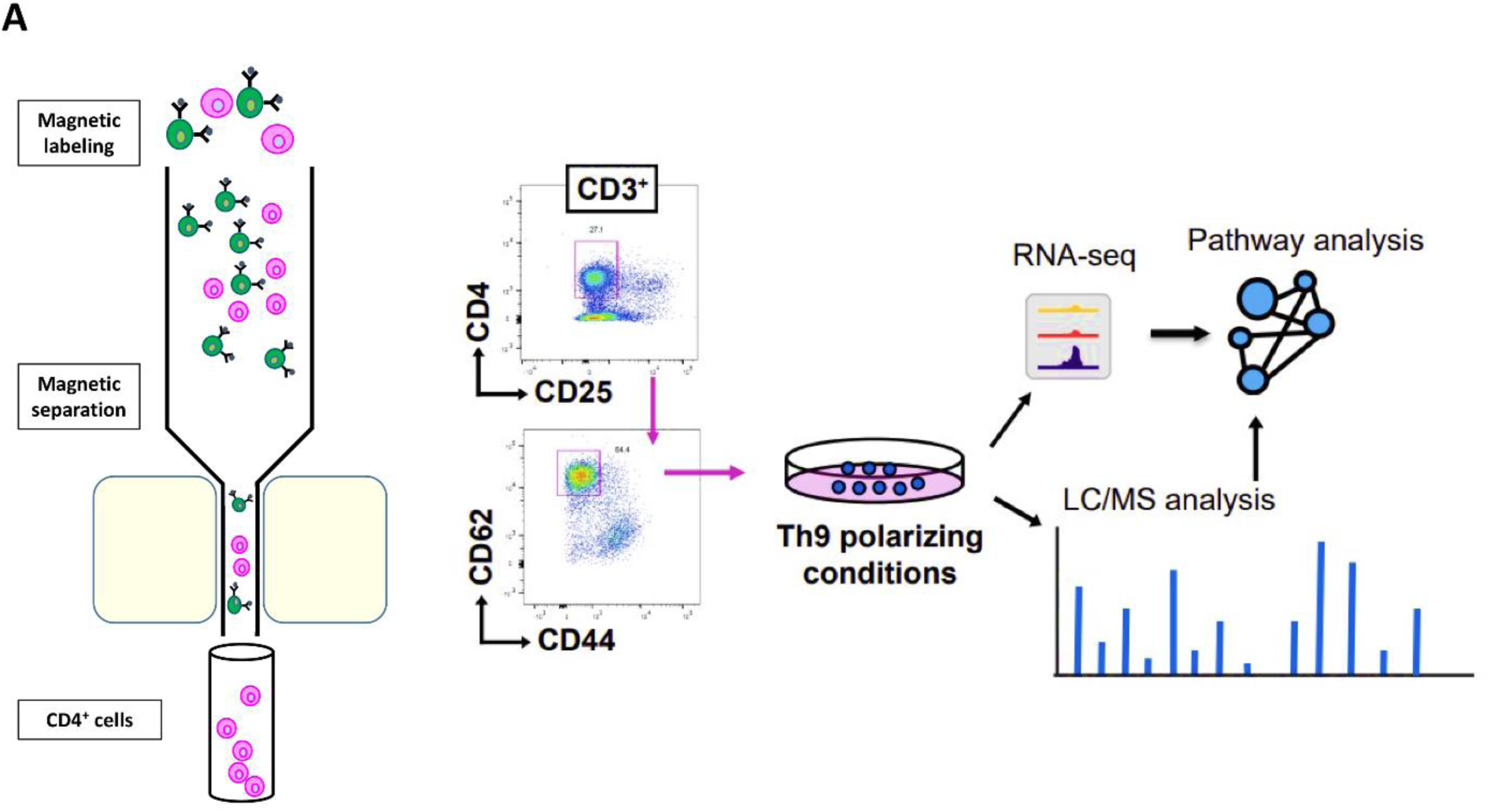
Th9 polarization from splenocytes. (**A**) Th9 cell workflow: Naïve CD3^+^CD4^+^CD62L^+^CD44^-^ cells enriched by magnetic column (MACS) and sorted from spleens of SAMP (WT) and DR3^-/-^xSAMP (KO) mice were differentiated into Th9 cells under Th9 polarizing conditions, and their molecular signature was investigated by RNA-seq and LC/MS-MS.

**Figure Suppl. 3:**
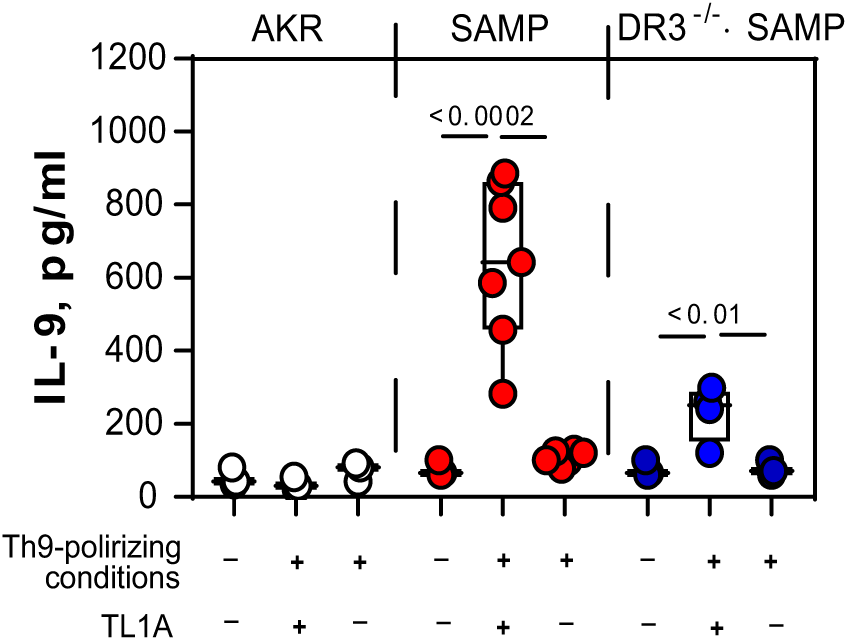
Functional DR3/TL1A system is essential for IL-9 protein secretion. **(A)** MACS sorted CD4^+^ T cells from 20-wks AKR, SAMP and DR3^-/-^×SAMP mice were stimulated under Th9 polarizing conditions with and without TL1A and IL-9 secretion was measured in cell supernatants by ELISA. Data, indicated as mean±SD, correspond to 2 independent experiments (n=7-12); statistical analysis was determined by 1-way Anova.

**Figure Suppl. 4:**
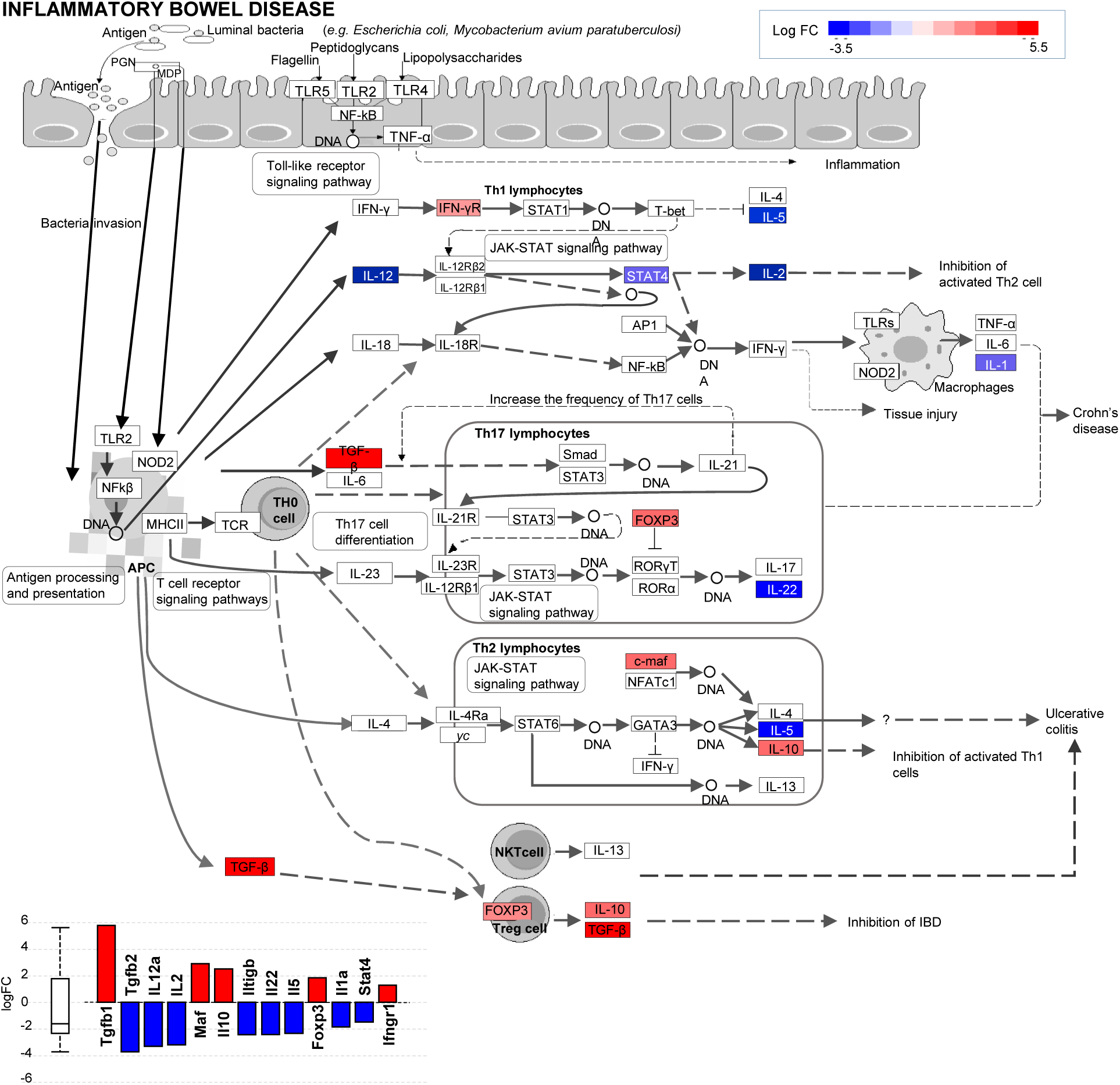
DR3 deletion determines changes at the transcriptome levels. (**A**) Inflammatory bowel disease (KEGG: 05321) pathway. The diagram is overlayed with the computed perturbation of each gene. The perturbation accounts both for the gene’s measured fold change and for the accumulated perturbation propagated from any upstream genes (accumulation). The highest negative perturbation is shown in dark blue, while the highest positive perturbation in dark red. The legend describes the values on the gradient; (**B**) Bar plot with all the differentially expressed genes in Inflammatory bowel disease (KEGG: 05321) are ranked based on their absolute value of log fold change. Upregulated genes are shown in red, downregulated genes are shown in blue. The box and whisker plot on the left summarizes the distribution of all the differentially expressed genes in the pathway. The box represents the 1st quartile, the median and the 3rd quartile, while the outliers are represented by circles. (n=5).

## References

1. Jergens, A.E., Parvinroo, S., Kopper, J. & Wannemuehler, M.J. Rules of Engagement: Epithelial-Microbe Interactions and Inflammatory Bowel Disease. Front Med (Lausanne) 8, 669913 (2021).

2. Yoo, J.H., Holubar, S. & Rieder, F. Fibrostenotic strictures in Crohn’s disease. Intest Res 18, 379–401 (2020).

3. Lewis, R.T. & Maron, D.J. Efficacy and complications of surgery for Crohn’s disease. Gastroenterol Hepatol (N Y) 6, 587–596 (2010).

4. Giuffrida, P., Corazza, G.R. & Di Sabatino, A. Old and New Lymphocyte Players in Inflammatory Bowel Disease. Dig Dis Sci 63, 277–288 (2018).

5. Vyas, S.P. & Goswami, R. A Decade of Th9 Cells: Role of Th9 Cells in Inflammatory Bowel Disease. Front Immunol 9, 1139 (2018).

6. Li, L. et al. Cytokine IL9 Triggers the Pathogenesis of Inflammatory Bowel Disease Through the miR21-CLDN8 Pathway. Inflamm Bowel Dis 24, 2211–2223 (2018).

7. Tian, L. et al. IL-9 promotes the pathogenesis of ulcerative colitis through STAT3/SOCS3 signaling. Biosci Rep 38 (2018).

8. Gerlach, K. et al. TH9 cells that express the transcription factor PU.1 drive T cell-mediated colitis via IL-9 receptor signaling in intestinal epithelial cells. Nat Immunol 15, 676–686 (2014).

9. Dardalhon, V. et al. IL-4 inhibits TGF-beta-induced Foxp3+ T cells and, together with TGF-beta, generates IL-9+ IL-10+ Foxp3(-) effector T cells. Nat Immunol 9, 1347–1355 (2008).

10. Stanko, K. et al. CD96 expression determines the inflammatory potential of IL-9-producing Th9 cells. Proc Natl Acad Sci U S A 115, E2940–E2949 (2018).

11. Li, Z. et al. Death Receptor 3 Signaling Controls the Balance between Regulatory and Effector Lymphocytes in SAMP1/YitFc Mice with Crohn’s Disease-Like Ileitis. Front Immunol 9, 362 (2018).

12. Bamias, G. et al. Expression, localization, and functional activity of TL1A, a novel Th1-polarizing cytokine in inflammatory bowel disease. J Immunol 171, 4868–4874 (2003).

13. Siakavellas, S.I. & Bamias, G. Tumor Necrosis Factor-like Cytokine TL1A and Its Receptors DR3 and DcR3: Important New Factors in Mucosal Homeostasis and Inflammation. Inflamm Bowel Dis 21, 2441–2452 (2015).

14. Bamias, G. et al. Role of TL1A and its receptor DR3 in two models of chronic murine ileitis. Proc Natl Acad Sci U S A 103, 8441–8446 (2006).

15. Butto, L.F. et al. Death-Domain-Receptor 3 Deletion Normalizes Inflammatory Gene Expression and Prevents Ileitis in Experimental Crohn’s Disease. Inflamm Bowel Dis 25, 14–26 (2019).

16. Danese, S. et al. Anti-TL1A Antibody PF-06480605 Safety and Efficacy for Ulcerative Colitis: A Phase 2a Single-Arm Study. Clin Gastroenterol Hepatol 19, 2324–2332 e2326 (2021).

17. Pizarro, T.T. et al. SAMP1/YitFc mouse strain: a spontaneous model of Crohn’s disease-like ileitis. Inflamm Bowel Dis 17, 2566–2584 (2011).

18. Lee, W.H. et al. BATF3 is sufficient for the induction of Il9 expression and can compensate for BATF during Th9 cell differentiation. Exp Mol Med 51, 1–12 (2019).

19. Tsuda, M. et al. A role for BATF3 in T(H)9 differentiation and T-cell-driven mucosal pathologies. Mucosal Immunol 12, 644–655 (2019).

20. Manocha, M. et al. Blocking CD27-CD70 costimulatory pathway suppresses experimental colitis. J Immunol 183, 270–276 (2009).

21. Souza, H.S., Elia, C.C., Spencer, J. & MacDonald, T.T. Expression of lymphocyte-endothelial receptor-ligand pairs, alpha4beta7/MAdCAM-1 and OX40/OX40 ligand in the colon and jejunum of patients with inflammatory bowel disease. Gut 45, 856–863 (1999).

22. Ahrens, R. et al. Intestinal macrophage/epithelial cell-derived CCL11/eotaxin-1 mediates eosinophil recruitment and function in pediatric ulcerative colitis. J Immunol 181, 7390–7399 (2008).

23. Heiseke, A.F. et al. CCL17 promotes intestinal inflammation in mice and counteracts regulatory T cell-mediated protection from colitis. Gastroenterology 142, 335–345 (2012).

24. Weder, B. et al. BCL-2 levels do not predict azathioprine treatment response in inflammatory bowel disease, but inhibition induces lymphocyte apoptosis and ameliorates colitis in mice. Clin Exp Immunol 193, 346–360 (2018).

25. Garcia, S. et al. Colony-stimulating factor (CSF) 1 receptor blockade reduces inflammation in human and murine models of rheumatoid arthritis. Arthritis Res Ther 18, 75 (2016).

26. Draghici, S. et al. A systems biology approach for pathway level analysis. Genome Res 17, 1537–1545 (2007).

27. Tarca, A.L. et al. A novel signaling pathway impact analysis. Bioinformatics 25, 75–82 (2009).

28. Bi, E. et al. Foxo1 and Foxp1 play opposing roles in regulating the differentiation and antitumor activity of TH9 cells programmed by IL-7. Sci Signal 10 (2017).

29. Macian, F. NFAT proteins: key regulators of T-cell development and function. Nat Rev Immunol 5, 472–484 (2005).

30. Brownlie, R. & Zamoyska, R. LAT polices T cell activation. Immunity 31, 174–176 (2009).

31. Maxwell, S., Chance, M.R. & Koyuturk, M. Linearity of network proximity measures: implications for set-based queries and significance testing. Bioinformatics 33, 1354–1361 (2017).

32. Yuan, A. et al. IL-9 antibody injection suppresses the inflammation in colitis mice. Biochem Biophys Res Commun 468, 921–926 (2015).

33. Shohan, M. et al. Th9 Cells: Probable players in ulcerative colitis pathogenesis. Int Rev Immunol 37, 192–205 (2018).

34. Rivera-Nieves, J. et al. Emergence of perianal fistulizing disease in the SAMP1/YitFc mouse, a spontaneous model of chronic ileitis. Gastroenterology 124, 972–982 (2003).

35. Richard, A.C. et al. The TNF-family ligand TL1A and its receptor DR3 promote T cell-mediated allergic immunopathology by enhancing differentiation and pathogenicity of IL-9-producing T cells. J Immunol 194, 3567–3582 (2015).

36. Meylan, F. et al. The TNF-family receptor DR3 is essential for diverse T cell-mediated inflammatory diseases. Immunity 29, 79–89 (2008).

37. Bamias, G. et al. Differential expression of the TL1A/DcR3 system of TNF/TNFR-like proteins in large vs. small intestinal Crohn’s disease. Dig Liver Dis 44, 30–36 (2012).

38. Weigmann, B. & Neurath, M.F. Th9 cells in inflammatory bowel diseases. Semin Immunopathol 39, 89–95 (2017).

39. Hufford, M.M. & Kaplan, M.H. A gut reaction to IL-9. Nat Immunol 15, 599–600 (2014).

40. Gerlach, K., McKenzie, A.N., Neurath, M.F. & Weigmann, B. IL-9 regulates intestinal barrier function in experimental T cell-mediated colitis. Tissue Barriers 3, e983777 (2015).

41. Chen, W. et al. Conversion of peripheral CD4+CD25-naive T cells to CD4+CD25+ regulatory T cells by TGF-beta induction of transcription factor Foxp3. J Exp Med 198, 1875–1886 (2003).

42. D’Ambrosio, D., Mariani, M., Panina-Bordignon, P. & Sinigaglia, F. Chemokines and their receptors guiding T lymphocyte recruitment in lung inflammation. Am J Respir Crit Care Med 164, 1266–1275 (2001).

43. Daig, R. et al. Increased interleukin 8 expression in the colon mucosa of patients with inflammatory bowel disease. Gut 38, 216–222 (1996).

44. Grimm, M.C., Elsbury, S.K., Pavli, P. & Doe, W.F. Interleukin 8: cells of origin in inflammatory bowel disease. Gut 38, 90–98 (1996).

45. Zhu, Y. et al. CXCL8 chemokine in ulcerative colitis. Biomed Pharmacother 138, 111427 (2021).

46. Masterson, J.C. et al. CCR3 Blockade Attenuates Eosinophilic Ileitis and Associated Remodeling. Am J Pathol 179, 2302–2314 (2011).

47. Colombel, J.F. et al. The safety of vedolizumab for ulcerative colitis and Crohn’s disease. Gut 66, 839–851 (2017).

48. Defendenti, C. et al. Significance of serum Il-9 levels in inflammatory bowel disease. Int J Immunopathol Pharmacol 28, 569–575 (2015).

49. Burns, R.C. et al. Antibody blockade of ICAM-1 and VCAM-1 ameliorates inflammation in the SAMP-1/Yit adoptive transfer model of Crohn’s disease in mice. Gastroenterology 121, 1428–1436 (2001).

50. Rodriguez-Palacios, A. et al. Stereomicroscopic 3D-pattern profiling of murine and human intestinal inflammation reveals unique structural phenotypes. Nat Commun 6, 7577 (2015).

51. Langmead, B. & Salzberg, S.L. Fast gapped-read alignment with Bowtie 2. Nat Methods 9, 357–359 (2012).

52. Li, B. & Dewey, C.N. RSEM: accurate transcript quantification from RNA-Seq data with or without a reference genome. BMC Bioinformatics 12, 323 (2011).

53. Kanehisa, M. & Goto, S. KEGG: kyoto encyclopedia of genes and genomes. Nucleic Acids Res 28, 27–30 (2000).

54. Kanehisa, M., Goto, S., Furumichi, M., Tanabe, M. & Hirakawa, M. KEGG for representation and analysis of molecular networks involving diseases and drugs. Nucleic Acids Res 38, D355–360 (2010).

55. Kanehisa, M. et al. Data, information, knowledge and principle: back to metabolism in KEGG. Nucleic Acids Res 42, D199–205 (2014).

56. Dobin, A. et al. STAR: ultrafast universal RNA-seq aligner. Bioinformatics 29, 15–21 (2013).

57. Trapnell, C. et al. Transcript assembly and quantification by RNA-Seq reveals unannotated transcripts and isoform switching during cell differentiation. Nat Biotechnol 28, 511–515 (2010).

