## Supplemental figures for "TL1A/DR3 signaling regulates the generation of pathogenic Th9 cells in experimental inflammatory bowel disease"

**Conflict of interest:** The authors have declared that they have no conflict of interest.

**Figure S. 1**

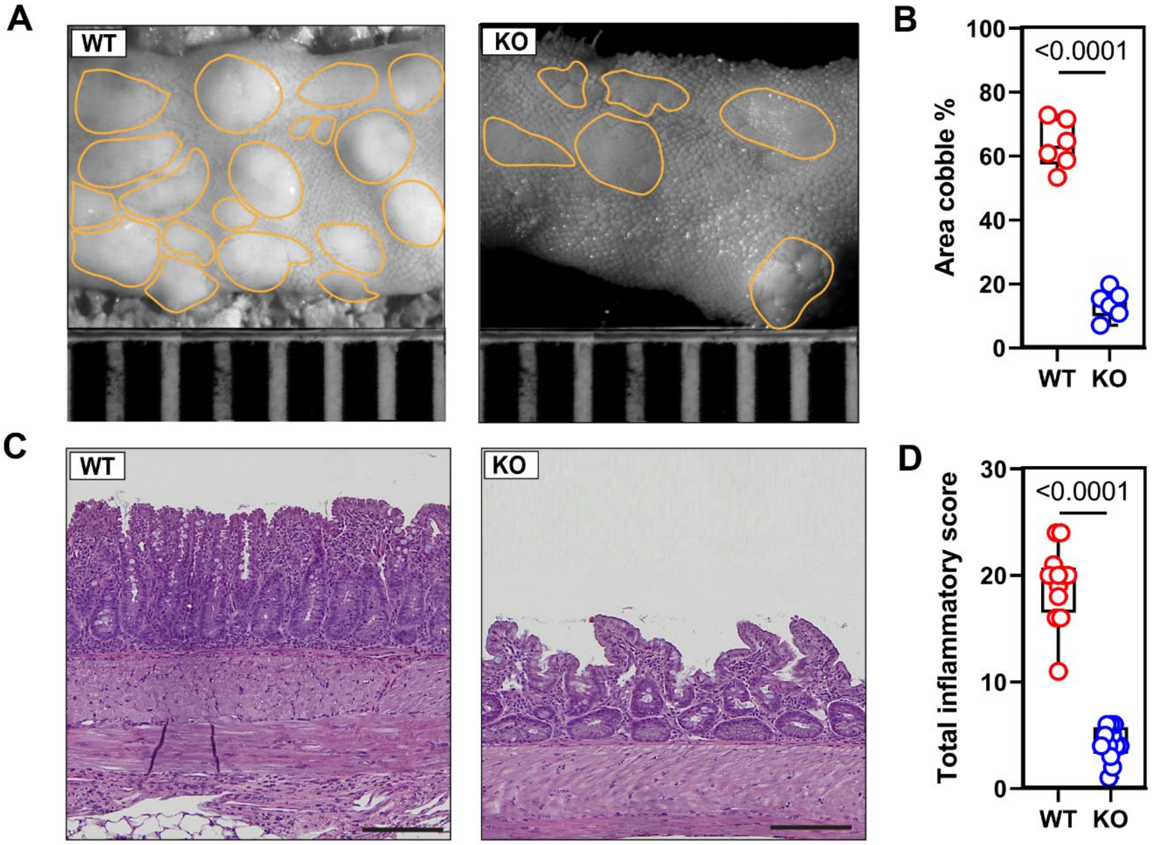


**Figure S. 1: DR3$^{-/-}$xSAMP (KO) mice exhibit a reduced inflammation compared to SAMP mice (WT).** (**A**) Mucosal architecture of fixed postmortem ileal specimens collected from SAMP (WT) and DR3$^{-/-}$xSAMP (KO) mice; (**B**) Cobblestone area expressed as percentage of total specimen calculated in the ileum of WT and KO mice (n=6). (**C**) Representative photomicrographs of ileal sections of 20-wk AKR and SAMP mice. Scale bar is 200 µm. (**D**) Total inflammatory score in 20-wks WT and KO mice (n=10).

**Figure S. 2**

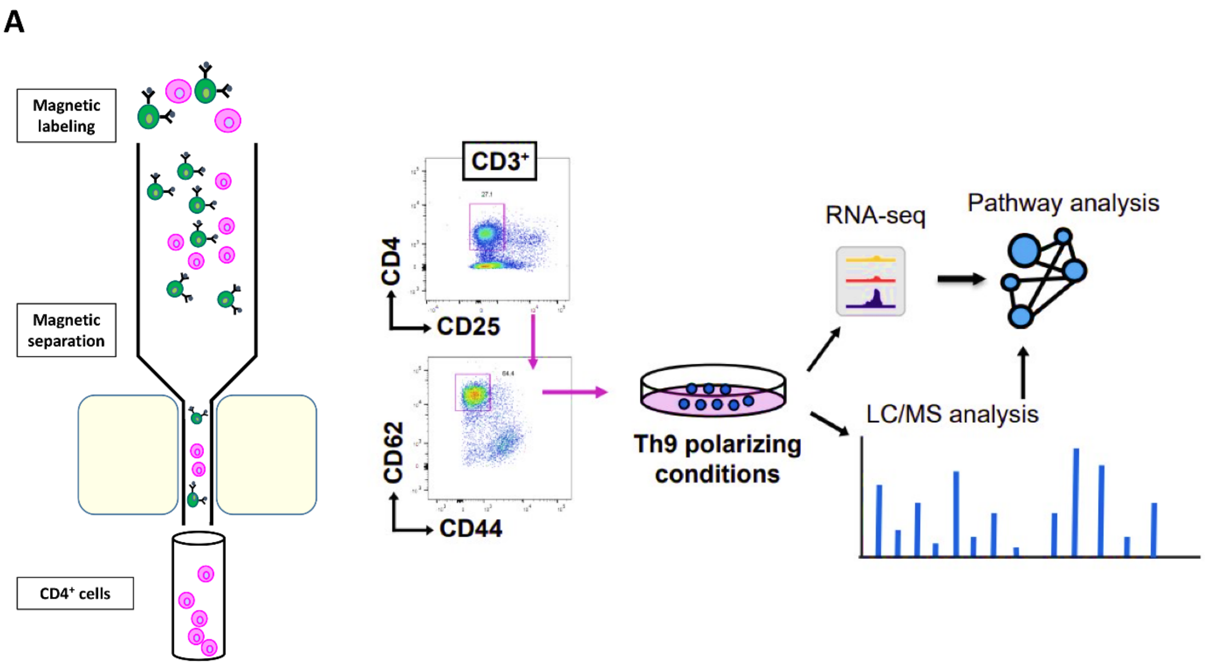


**Figure S. 2: Th9 polarization from splenocytes.** (**A**) Th9 cell workflow: Naïve $CD3^{+}CD4^{+}CD62L^{+}CD44^{-}$ cells enriched by magnetic column (MACS) and sorted from spleens of SAMP (WT) and $DR3^{-/-}$xSAMP (KO) mice were differentiated into Th9 cells under Th9 polarizing conditions, and their molecular signature was investigated by RNA-seq and LC/MS-MS.

**Figure S.3**

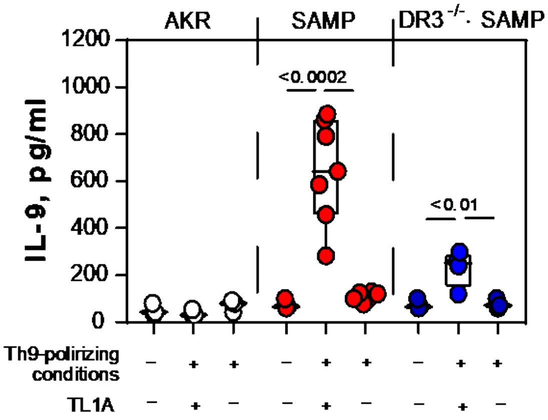


**Figure S. 3: Functional DR3/TL1A system is essential for IL-9 protein secretion.** (**A**) MACS sorted CD4+ T cells from 20-wks AKR, SAMP and DR3$^{-/-}$×SAMP mice were stimulated under Th9 polarizing conditions with and without TL1A and IL-9 secretion was measured in cell supernatants by ELISA. Data, indicated as mean±SD, correspond to 2 independent experiments (n=7-12); statistical analysis was determined by 1-way Anova.

**Figure S. 4**

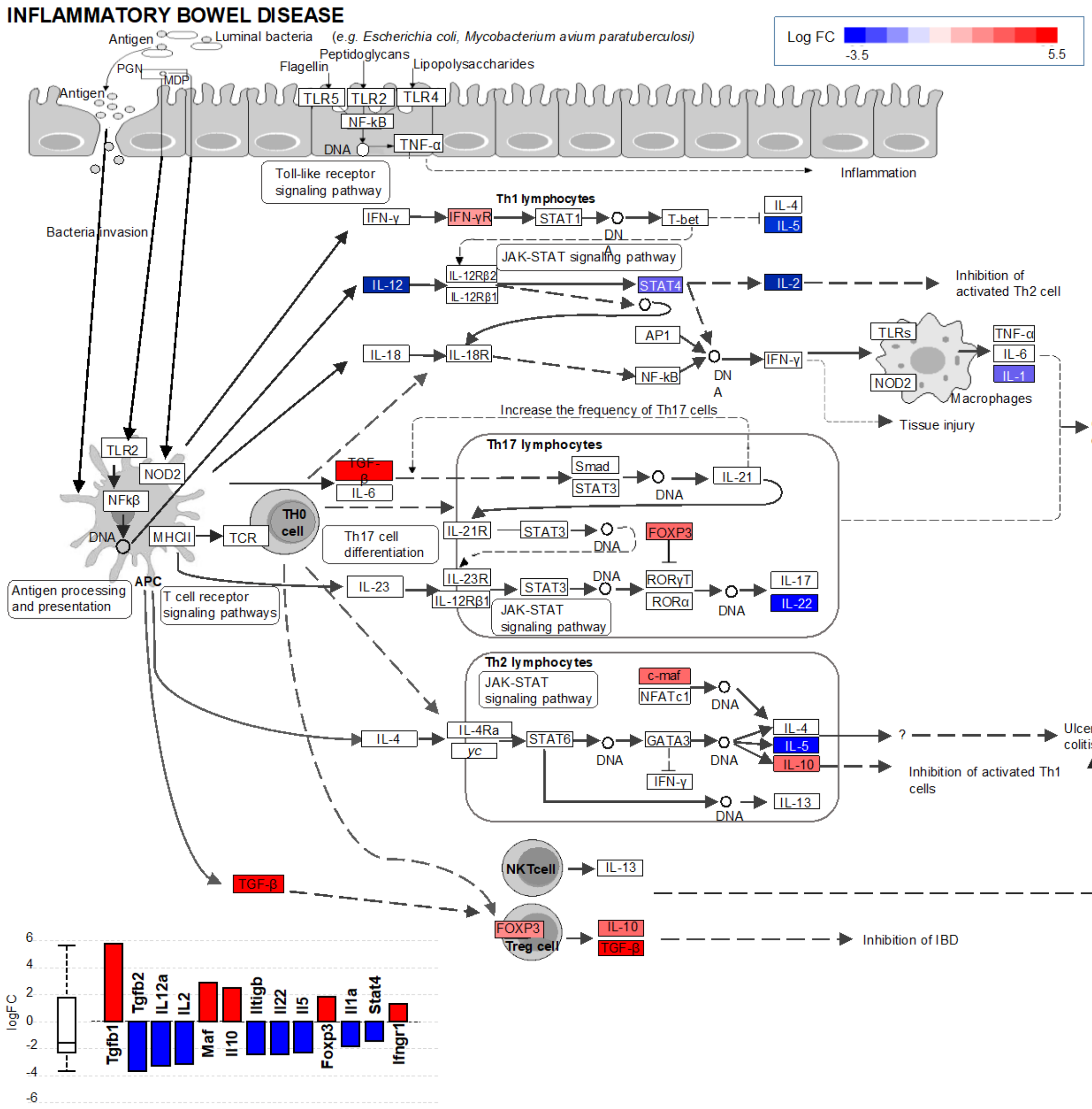


**Figure S. 4: DR3 deletion determines changes at the transcriptome levels.** (**A**) Inflammatory bowel disease (KEGG: 05321) pathway. The diagram is overlayed with the computed perturbation of each gene. The perturbation accounts both for the gene's measured fold change and for the accumulated perturbation propagated from any upstream genes (accumulation). The highest negative perturbation is shown in dark blue, while the highest positive perturbation in dark red. The legend describes the values on the gradient; (**B**) Bar plot with all the differentially expressed genes in Inflammatory bowel disease (KEGG: 05321) are ranked based on their absolute value of log fold change. Upregulated genes are shown in red, downregulated genes are shown in blue. The box and whisker plot on the left summarizes the distribution of all the differentially expressed genes in the pathway. The box represents the 1st quartile, the median and the 3rd quartile, while the outliers are represented by circles. (n=5).
